# Conservation of *N*-hydroxy-pipecolic acid-mediated systemic acquired resistance in crop plants

**DOI:** 10.1101/537597

**Authors:** Eric C. Holmes, Yun-Chu Chen, Elizabeth Sattely, Mary Beth Mudgett

**Affiliations:** Department of Chemical Engineering, Stanford University, Stanford, CA, 94305, USA; Department of Biology, Stanford University, Stanford, CA, 94305, USA; Howard Hughes Medical Institute

**Keywords:** Systemic acquired resistance, plant innate immunity, *N*-hydroxy-pipecolic acid, biosynthetic pathway reconstitution

## Abstract

Signal propagation and the coordination of whole-organism responses in plants rely heavily on small molecules. Systemic acquired resistance (SAR) is one such process in which long-distance signaling activates immune responses in uninfected tissue as a way to limit the spread of a primary, localized infection. Recently, *N*-hydroxy pipecolic acid (NHP) was discovered and shown to coordinate SAR in *Arabidopsis*. Here, we provide metabolic and biochemical evidence that NHP is conserved across the plant kingdom and demonstrate a role for NHP in mediating SAR responses in tomato and pepper. We reconstituted the NHP biosynthetic pathway *in planta* and show that transient expression of two NHP biosynthetic genes in tomato induces enhanced resistance to a bacterial pathogen in distal tissue. Our results suggest engineering strategies to induce NHP-mediated SAR are a promising route to improve broad-spectrum pathogen resistance in crops.

**IN BRIEF:** Engineering NHP production is a promising strategy to enhance disease resistance in crops.

**HIGHLIGHTS:**

- *Arabidopsis N*-hydroxy-pipecolic acid (NHP) pathway is conserved across the plant kingdom
- Application of NHP to tomato and pepper plants induces a robust SAR response
- Metabolic engineering of the *Arabidopsis* NHP pathway in *Solanum lycopersicum* leads to enhanced NHP production and defense priming
- Genetic engineering for enhanced NHP production is a promising strategy to protect crop plants from multiple pathogens

## INTRODUCTION

Plants resist infection using a sophisticated innate immune system to detect and respond to pathogens in their environment (Spoel and Dong, 2012). Pathogens are first detected by cell surface pattern recognition receptors that recognize conserved microbial associated molecular patterns or MAMPs (Couto and Zipfel, 2016). Initial pathogen detection can be augmented by intracellular recognition of specific pathogen-derived effector proteins by nucleotide-binding domain and leucine rich repeat containing (NLR) proteins (Cui et al., 2015). This combined detection system is inherently flexible, allowing for surveillance of multiple pathogens and the evolution of disease resistance specificity across the plant kingdom. Inputs from these receptors converge on central, well-conserved signal transduction networks that use ubiquitous hormones such as salicylic acid (SA), jasmonic acid (JA), and ethylene (ET) (Glazebrook, 2005) to activate and amplify diverse defense responses. The multiple outputs of innate immune signaling pathways can vary by plant species and include species-specific biosynthesis of antimicrobial metabolites (Ahuja et al., 2012). Integration of conserved signaling pathways with species-specific defense mechanisms is central for mounting an effective plant immune response (Pieterse et al., 2009). The modular nature of the plant immune system has enabled the engineering of desired resistance traits in different plant species. For example, transferring the unique pathogen-detection capabilities from one plant family to others by genetic engineering has led to lines with broad spectrum and enhanced disease resistance relative to wild-type plants (Rodriguez-Moreno et al., 2017).

In addition to local immune responses, plants utilize a circulating, chemical immune system to provide protection throughout the plant body. One classic example is systemic acquired resistance (SAR). During SAR, pathogen infection of primary (local) plant tissue leads to the production of mobile plant signals that move long-distances to activate immune responses in secondary (distal) uninfected tissues. Such defense priming can prevent the spread of the initial infection or the occurrence of a new infection, providing resistance to a broad spectrum of potential pathogens (Fu and Dong, 2013). In some cases, SAR can be transferred to the plant progeny in next generation via epigenetic mechanisms (Ton, 2012). Elucidating the molecular mechanisms that govern long-distance defense priming is an intense area of research given that SAR appears to be conserved in plant families across the plant kingdom (Shah and Zeier, 2013) and there is potential for enhancing plant protection via engineering strategies.

Several molecules shown to be involved in long-distance signal transduction for SAR, including SA (Klessig et al., 2018), methyl salicylate (MeSA) (Park et al., 2007), azelaic acid (AzA) (Jung et al., 2009), glycerol-3-phosphate (G3P) (Chanda et al., 2011), dehydroabietinal (DA) (Chaturvedi et al., 2012), and pipecolic acid (Pip) (Navarova et al., 2012). Among these metabolites, *N*-hydroxy pipecolic acid (NHP) has recently emerged as a key player that is required to initiate SAR (Chen et al., 2018; Hartmann et al., 2018). Notably, direct treatment of wild-type *Arabidopsis thaliana* leaves with NHP makes distal, untreated leaves more resistant to pathogen growth, with levels of protection comparable to that observed in a classical SAR response initiated by a primary infection. These SAR responses are attenuated in SA biosynthetic mutants, highlighting the importance of SA in NHP-induced SAR signaling (Hartmann et al., 2018). Furthermore, NHP provides protection against the oomycete pathogen *Hyalopersonospora arabidopsisdis* (Hartmann et al., 2018) and the bacterial pathogen *Pseudomonas syringae* (Chen et al., 2018; Hartmann et al., 2018), indicating that NHP signaling results in broad-spectrum disease resistance.

The discovery of NHP as a central signaling molecule in SAR provides the basis for elucidating how species-specific pathogen detection and defense output responses integrate with core signaling pathways conserved in the plant immune system. A critical first step is to determine the presence and role of NHP in plants other than *Arabidopsis*. NHP has been shown to be derived from lysine, and previous work has associated three enzymes with the NHP biosynthetic pathway: (1) the aminotransferase ALD1 (AGD2-LIKE DEFENSE RESPONSE PROTEIN 1; (Navarova et al., 2012)), which generates an α-keto acid from lysine; (2) a reductase, one of which is SARD4 (SAR-DEFICIENT 4; (Ding et al., 2016)), to generate Pip; and (3) FLAVIN-DEPENDENT MONOOXYGENASE 1 (FMO1; (Chen et al., 2018; Hartmann et al., 2018)), an enzyme that installs the hydroxyl amine of NHP and is known to be a critical regulator of SAR (Bartsch et al., 2006; Koch et al., 2006; Mishina and Zeier, 2006) (Figure 1A). This collection of biosynthetic genes is an ideal starting point to determine the conservation of NHP across the plant kingdom.

**Figure 1.**
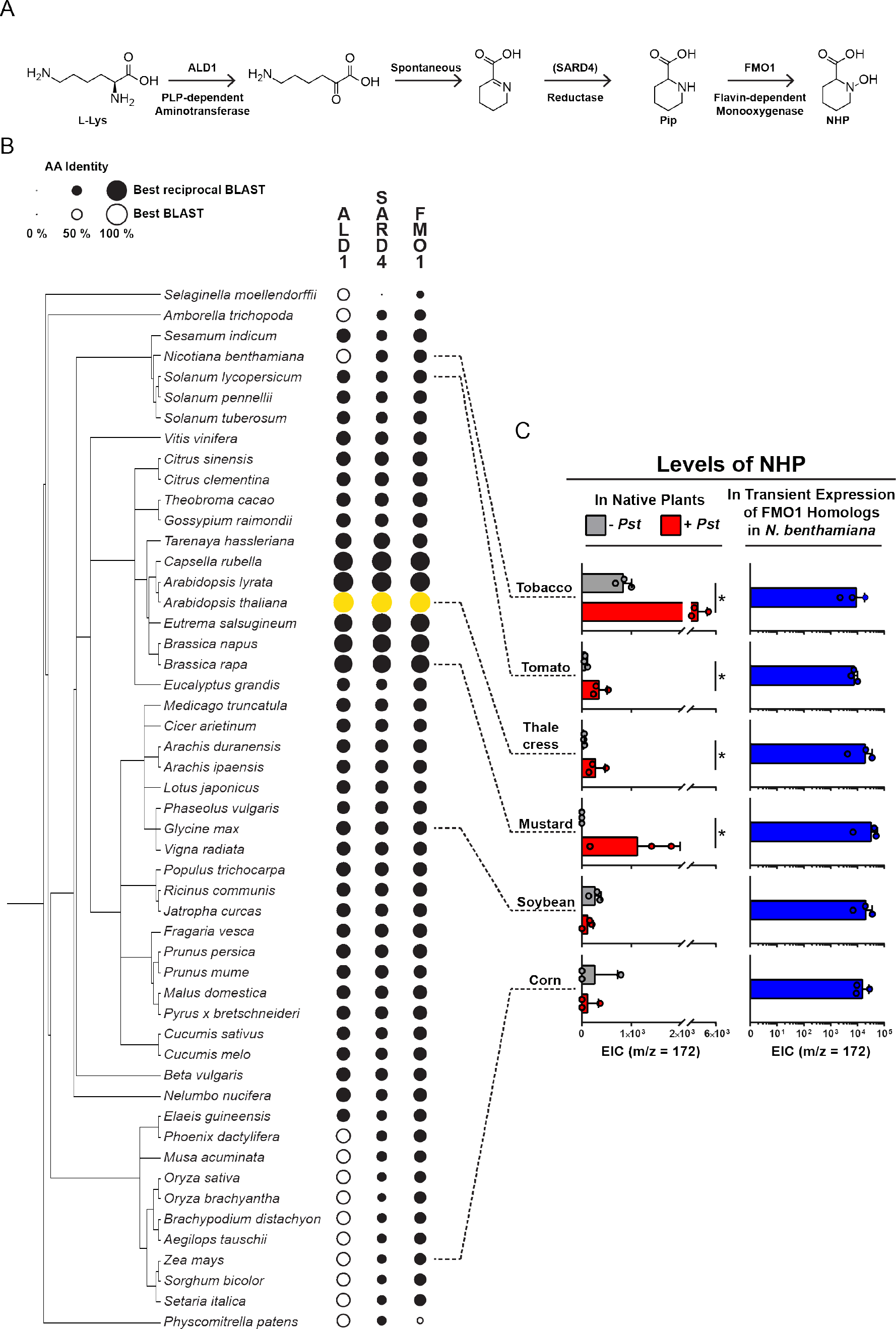
Evidence of NHP production across the plant phylogeny. (A) Canonical biosynthetic pathway to *N*-hydroxypipecolic acid (NHP). (B) Phylogenetic tree of sequenced plant genomes created using the PhyloT tree generator (http://phylot.biobyte.de/index.html). Circles are scaled by the percent amino acid identity of the best BLAST (blastp) hit between the *Arabidopsis thaliana* NHP biosynthetic proteins (yellow) and the ALD1, SARD4, and FMO1 homologs of the respective plant proteome (black). Solid circles signify that the homolog returned the respective *A. thaliana* protein in a reverse-BLAST into the *A. thaliana* proteome (best reciprocal BLAST). (C) Evidence for NHP production in six plant species. Dotted lines label the species and common name, respectively, for plants examined: *N. benthamiana*, tobacco; *S. lycopersicum*, tomato; *A. thaliana*, thale cress; *B. rapa*, mustard; *Glycine max*, soybean; *Zea mays*, corn. To measure endogenous NHP, seedlings were grown hydroponically and then mock-treated (− *Pst*; grey) or elicited with *Pseudomonas syringae* pathovar *tomato* strain DC3000 (+ *Pst*; red) for 48 hpi. NHP was detected using GC-MS. Bars indicate extracted ion abundance (EIC) for NHP (m/z = 172). Asterisks indicate a significant NHP increase in *Pst*-elicited plants (one-tailed t-test; * *P* < 0.05). Pip and NHP-Glc levels for this experiment are shown in Figure S2. To measure enzyme activity, FMO1 homologs from each plant with highest identify with *Arabidopsis* FMO1 were cloned into *Agrobacterium tumefaciens* and then transiently expressed in *N. benthamiana* leaves in the presence of 1 mM Pip. Bars (blue) indicate EIC of NHP measured in leaves expressing the respective FMO1 homolog. Pip levels and controls for this experiment are graphed in Figure S3. Bars represent the mean ± STD of three biological replicates. Values reported as zero indicate no detection of metabolites.

Here we show that FMO1 homologs from phylogenetically distant species can catalyze the conversion of Pip to NHP using transient expression in *Nicotiana benthamiana*. We also report the detection of NHP in a variety of monocot and dicot seedling extracts, and the inducible accumulation of NHP in plant species in the Brassicaceae and Solanaceae in response to the bacterial pathogen *Pseudomonas syringae*. Furthermore, we demonstrate that application of synthetic NHP induces SAR in *Solanum lycopersicum* (tomato) and *Capsicum annuum* (pepper), two representative model vegetable crop plants, protecting distal leaves from bacterial infection. Finally, we show that NHP biosynthesized in tomato leaves using *Agrobacterium*-mediated transient expression of just two of the NHP biosynthetic pathway genes results in SAR in distal tissue. These results demonstrate that NHP is not only synthesized in plant species outside the Brassicaceae, but also provides SAR protection. Moreover, our work indicates that metabolic engineering of the NHP pathway is a promising approach to override and perhaps optimize innate immune responses in important crop plants.

## RESULTS

### Conservation of NHP biosynthetic enzymes in plant kingdom

As a first step to determine whether NHP may function as a SAR signal throughout the plant kingdom, we investigated whether the biosynthetic pathway is conserved (Figure 1). We used NCBI’s protein BLAST tool (https://blast.ncbi.nlm.nih.gov/Blast.cgi) to compare the percent amino acid identity between the three proteins associated with NHP biosynthesis in *Arabidopsis:* ALD1, SARD4 and FMO1 (Figure 1A) and their closest homologs in 50 other plant species. We then performed a reverse-BLAST of each homolog back to the *Arabidopsis* proteome to identify reciprocal-BLAST pairs, a common criteria for determining orthologous proteins (Moreno-Hagelsieb and Latimer, 2008). We found that 37 of the 50 plant species analyzed contain an entire orthologous NHP biosynthetic pathway (ALD1, SARD4, and FMO1) and that all but one (the moss, *Physcomitrella patens*) harbors an orthologous FMO1 protein (Figure 1B).

To understand how the apparent NHP pathway conservation compares to other signaling and defense metabolites, we performed a similar analysis using the biosynthetic enzymes for salicylic acid (SA) and jasmonic acid (JA), two widely conserved defense hormones, and indole glucosinolate (GSL), a Brassicaceae-specific defense metabolite. Specifically, we analyzed isochorismate synthase 1 (Wildermuth et al., 2001) for SA, OPDA reductase (Stintzi and Browse, 2000) and jasmonate resistance 1 (Staswick and Tiryaki, 2004) for JA, and S-alkyl-thiohydroximate lyase (Mikkelsen et al., 2004) and cytochrome P450 monooxygenase CYP83B1 (Bak et al., 2001) for GSL and compared the profiles of homologous proteins across the plant kingdom with those in the NHP pathway (Figure S1). The pattern and relative degree of sequence similarity of best reciprocal-BLAST orthologs in the NHP pathway corresponds with the pathway patterns for SA and JA, but differs from the Brassicaceae-specific glucosinolate pathway (Figure S1). This analysis suggests that the NHP biosynthetic pathway is well conserved and that there is a tight correlation between NHP and SA biosynthesis amongst plant species. NHP signaling is known to initiate SA biosynthesis in *Arabidopsis* (Chen et al., 2018), highlighting a direct link between these two metabolites in the establishment of SAR.

### NHP production across plant kingdom

The conservation of the NHP biosynthetic pathway (Figure 1B) suggests that NHP may be produced in species throughout the plant kingdom. To examine this possibility, we compared the level of endogenous and pathogen-elicited NHP pathway metabolites in *Arabidopsis* with that of four dicot plants (*Brassica rapa* (field mustard), Order Brassicales; *N. benthamiana* and *S. lycopersicum* (tomato), Order Solanales; and *Glycine max* (soybean), Order Fabales) and one monocot plant (*Zea mays* (corn), Order Poaceae) (Figure 1C). Seedlings were grown axenically in hydroponic media for 1 to 2 weeks and then elicited for 48 h with the bacterial pathogen *Pseudomonas syringae* pathovar *tomato* strain DC3000 (*Pst*). Liquid chromatography and gas chromatography followed by mass spectrometry (LC-MS and GC-MS) were used to quantify endogenous and *Pst-*elicited biosynthesis of Pip, NHP and NHP-glucoside (NHP-Glc), a glycosylated conjugate of NHP (Chen et al., 2018).

NHP was detected in untreated seedlings of *N. benthamiana*, tomato, soybean, and corn, but not *B. rapa* (Figure 1C) and levels increased in seedlings of *N. benthamiana*, tomato, and *B. rapa* after *Pst* elicitation (Figure 1C). By contrast, NHP levels in *Pst-*treated seedlings of soybean and corn were similar to that of the untreated seedlings (Figure 1C). Levels of Pip, the precursor of NHP, also increased in response to *Pst* treatment (Figure S2). Notably, NHP-Glc only accumulated in *Arabidopsis* and *B. rapa* (Figure S2). These data reveal that in addition to *Arabidopsis*, five diverse species produce NHP. They also provide evidence that increased accumulation of NHP in tissues in response to pathogen-elicitation outside of the Brassicaceae, while the production of NHP-Glc may be unique to the Brassicaceae (Figure S2).

### Biochemical activity of putative FMO1 orthologs

Next we analyzed the biochemical activity of the putative FMO1 orthologs identified for the five plant species examined in Figure 1C. We cloned each *FMO1-*like gene (BraA06g014860 from *B.* rapa; Glyma13g17340 from *G. max*; Solyc07g042430 from *S. lycopersicum*; Niben101Scf05682g00009 from *N. benthamiana*; AC191071.3_FGP001 from *Zea mays*) and transiently expressed each in *N. benthamiana* leaves using *Agrobacterium tumefaciens* strains. *Arabidopsis FMO1* was used as a positive control. Leaves were also infiltrated with 1 mM Pip to provide ample substrate for the reaction. Expression of each putative FMO1 ortholog led to the production of NHP (Figure 1C and S3). NHP was the major metabolite identified by GC-MS for all plant FMO1-like enzymes tested and no differences in metabolites profiles were apparent in the respective leaf extracts in comparison to those expressing *Arabidopsis* FMO1. Leaves expressing GFP (negative control) or an inactive *Arabidopsis* FMO1 mutant (FMO1 G17A/G19A) (Chen et al., 2018) did not produce any NHP (Figure S3). These data show that *FMO1-*like genes from diverse plant species encode orthologous proteins that can catalyze the *N*-hydroxylation of Pip when expressed in a heterologous plant, providing biochemical evidence that monooxygenases capable of producing NHP are conserved across the plant kingdom.

### Exogenous application of NHP to Solanaceous plants induces SAR

Previously, we demonstrated that exogenous application of purified NHP to the roots or leaves of *Arabidopsis* plants was sufficient to induce SAR signaling and inhibit the growth of a virulent bacterial strain, *Pseudomonas syringae* pathovar maculicola *ES4326* (Chen et al., 2018). The presence of FMO1-like enzymes and *Pst-*elicited production of NHP in tomato and *N. benthamiana* (Figure 1) suggested that NHP might also function as a SAR signal within the Solanaceae. To test this, we assessed the bioactivity of purified NHP in fully expanded leaves of a 4 to 5-week old tomato plant (*Solanum lycopersicum* cultivar VF36). Solutions of 10 mM MgCl_2_ (Mock) or 10 mM MgCl_2_ containing 1 mM NHP were infiltrated into the two leaflets closest to the main stem (referred to as the “top” leaflets) (Figure 2A). 24 h later, the remaining three leaflets (referred to as the “bottom” leaflets) were challenged with a 1×10^5^ cfu/ml suspension of *Pst*, a virulent strain that infects VF36 tomato plants (Figure 2A). At 4 days post-inoculation (dpi), the titer of *Pst* in the bottom leaflets was quantified (Figure 2B) and leaf symptoms were photographed (Figure 2C). The titer of *Pst* was significantly lower in the bottom leaflets of leaves that were treated with 1 mM NHP compared to Mock treated leaves (Figure 2B). In addition, disease symptoms (i.e. bacterial speck, leaf yellowing and leaf necrosis) were much less in the *Pst* infected bottom leaflets of NHP-treated leaves compared to those of Mock-treated leaves (Figure 2C). These data show that local treatment of NHP in top leaflets is sufficient to induce disease resistance in neighboring bottom leaflets of the same leaf, providing evidence that NHP is a bioactive SAR signal in tomato.

**Figure 2.**
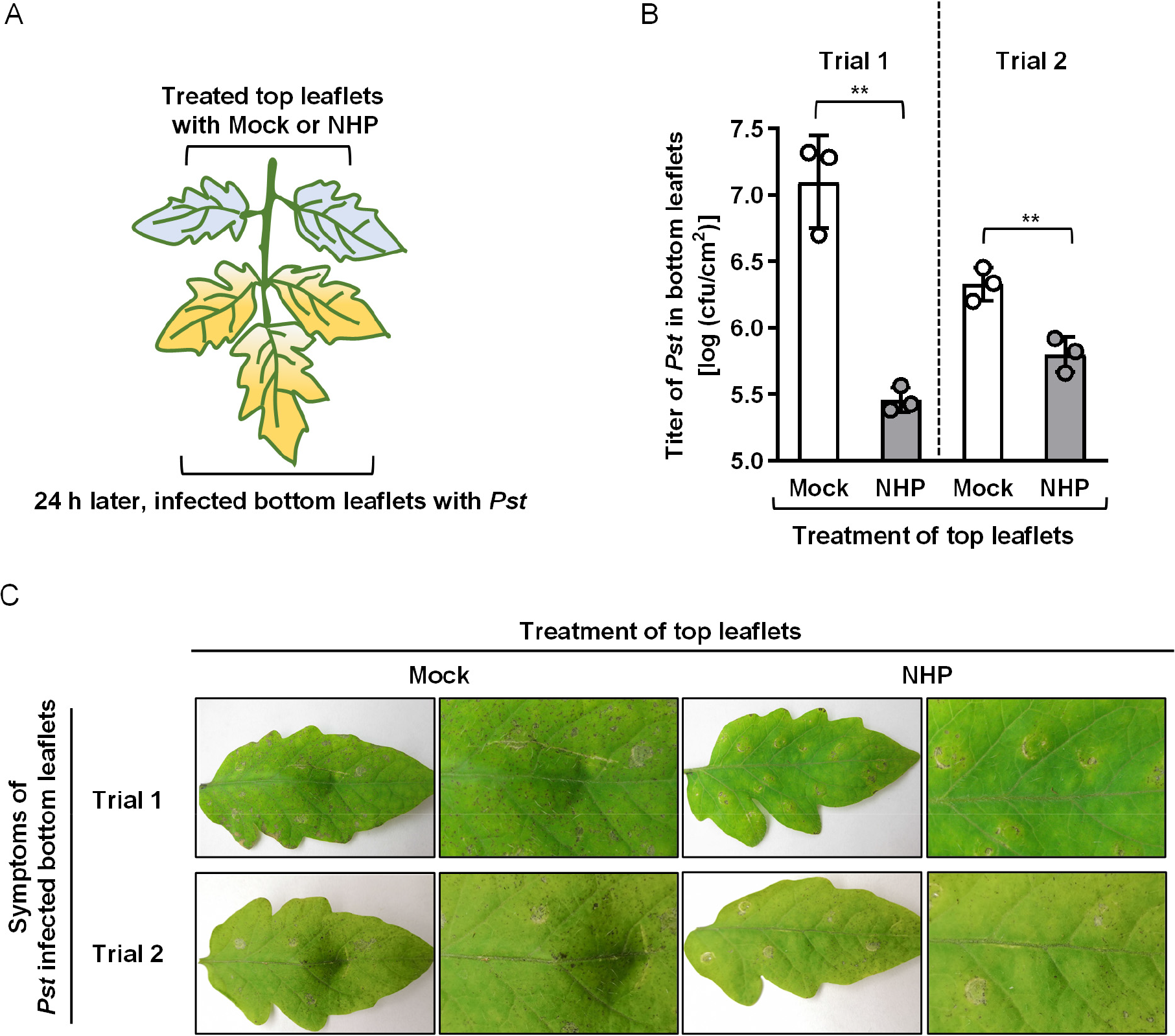
Local treatment of NHP induces systemic defense in tomato, *Solanum lycopersicum* cultivar VF36, leaves. (A) Diagram showing treatment design of tomato leaves for SAR experiments. Two top leaflets of a tomato leaf were infiltrated with 10 mM MgCl_2_ (mock) or 10 mM MgCl_2_ containing 1 mM NHP. After chemical incubation for 24 h, three bottom leaflets of the same layer were inoculated with a 1×10^5^ cfu/ml suspension of *Pst*. (B) Titer of *Pst* in bottom leaflets at 4 dpi. Bars represent mean ± STD of three leaflets from one representative plant in two independent trials. Asterisks denote the significant differences between indicated samples using a one-tailed t-test (** *P* < 0.01). (C) Disease symptoms of bottom leaflets infected with *Pst* at 4 dpi. For each treatment, panels show symptoms of a whole leaflet (left) and an enlarged region of the respective leaflet (right). The experiment was repeated for three times, and similar results were obtained.

To extend our analysis to an additional crop plant, we tested the bioactivity of NHP in pepper plants, *Capsicum annuum*. We infiltrated two lower, fully expanded leaves of a 4 to 5-week old pepper plant (cultivar Early Calwonder) with 10 mM MgCl_2_ (Mock) or 10 mM MgCl_2_ containing 2 mM NHP. Twenty-four h later, an upper leaf was inoculated with a 1×10^4^ cfu/ml suspension of *Xanthomonas euvesicatoria* strain 85-10 (*Xe 85-10*), a virulent strain that infects Early Calwonder pepper plants (Figure S4A). At 10 dpi, there was a significant decrease in *Xe 85-10* growth (Figure S4B) and symptom development (Figure S4C) in pepper plants treated with NHP compared to Mock. These results demonstrate that NHP treatment of pepper plants is sufficient to prime a systemic defense response. Moreover, in combination with our results in tomato, these data provide strong evidence that component(s) that detect NHP and activate SAR are present in the Solanaceae.

### Reconstitution of Arabidopsis NHP biosynthetic pathway in *N. benthamiana*

We hypothesized that reconstitution of NHP biosynthesis de novo from lysine would require expression of all genes associated with the *Arabidopsis* NHP pathway (ALD1/SARD4/FMO1). In preliminary experiments, we co-infiltrated a mixture of three *Agrobacteria* strains each harboring one of the three *Arabidopsis* NHP pathway genes into leaves of *N. benthamiana* and observed significant accumulation of NHP. The levels of NHP in this experiment were 10-100 fold higher than levels observed after expression of FMO1 alone with 1 mM Pip supplementation (Figures 3 and S3). This suggests that the substrate Pip is limiting when it is infiltrated as a solution into leaves. Co-expression of FMO1 with enzymes that direct the conversion of Lys to Pip alleviates this substrate availability problem. Subsequent optimization revealed SARD4 was not necessary for NHP production in *N. benthamiana* leaves, as NHP accumulation was similar in leaves expressing ALD1/SARD4/FMO1 and ALD1/FMO1 (Figure 3). This result suggests that *N. benthamiana* leaves provide sufficient endogenous reductase activity for conversion of dehydro-Pip to Pip. Collectively, these data show that the full *Arabidopsis* NHP pathway can be reconstituted in a heterologous system. Furthermore, utilizing the endogenous pool of Lys for NHP biosynthesis can lead to high levels of NHP production in the absence of pathogen elicitation.

**Figure 3.**
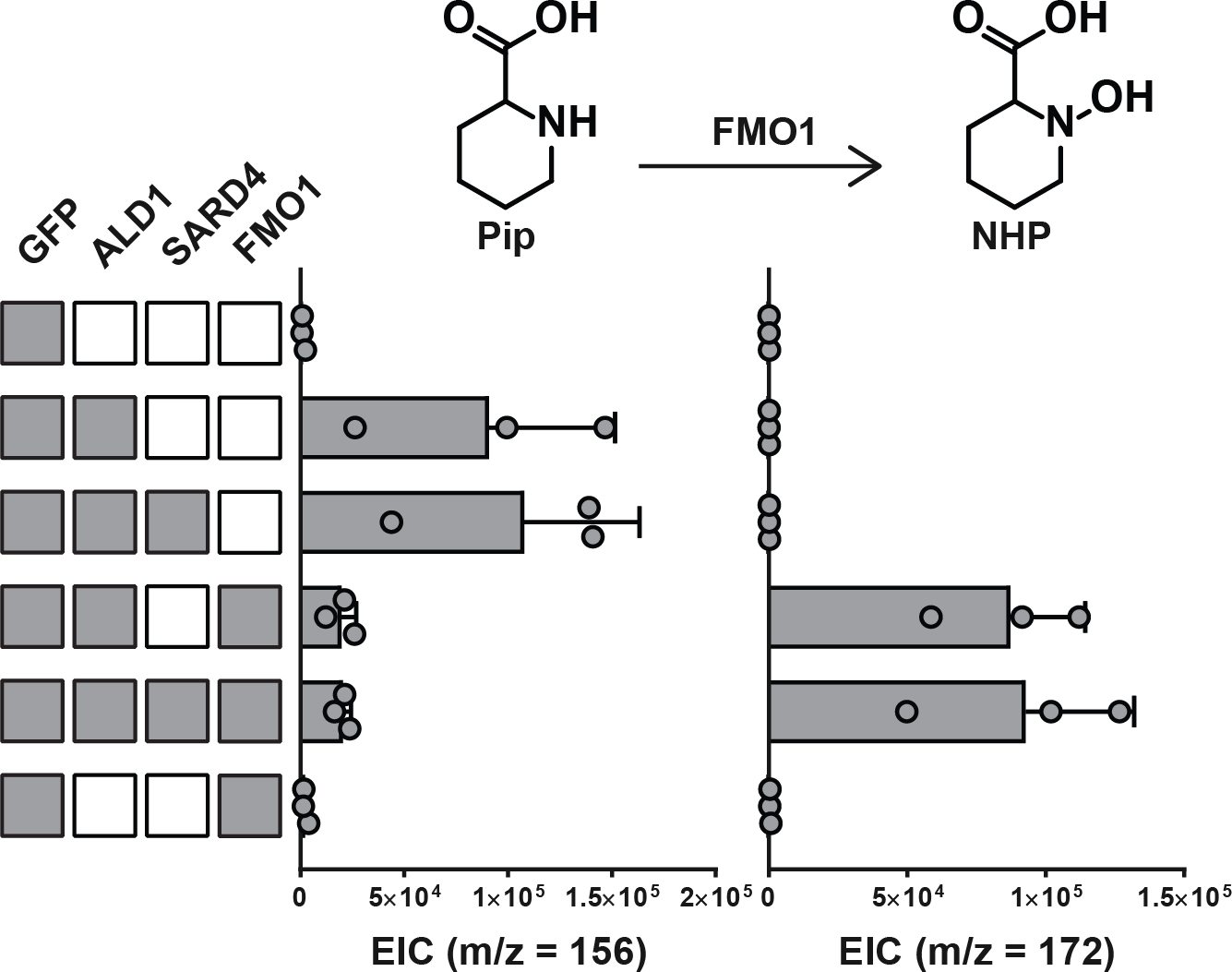
Expression of NHP pathway enzymes in *N. benthamiana*. Ion abundances of Pip (m/z = 156) and NHP (m/z = 172) in *N. benthamiana* leaves expressing combinations of *Arabidopsis ALD1*/*SARD4*/*FMO1* and *GFP* genes for 24 h determined by GC-MS. Each row represents a different enzyme set where a grey box indicates that an *Agrobacterium* strain containing that gene was included in the infiltration. For each biosynthetic gene, an *Agrobacterium* strain was infiltrated at an OD_600_ of 0.1 and a strain harboring *GFP* was included to normalize the total OD_600_ of infiltration at 0.4. Bars represent the mean ± STD of three biological replicates. Levels reported as zero indicate no detection of metabolites.

In addition to NHP, we also noted a collection of additional mass features in LC-MS and GC-MS chromatograms that are co-produced with NHP in *N. benthamiana* leaves expressing ALD1 and FMO1. Careful investigation of each of the most abundant mass signatures revealed two sets of compounds. The first group is proposed to be structurally related to NHP based on MS-MS analysis (Figures S5, compounds 1-4). Several of these (compounds 1-3) also accumulated when an authentic synthetic standard of NHP was heated, suggesting that they may be degradation products and potentially formed during the leaf extraction process. The second group is proposed to be derived from proline and was dependent upon FMO1, but not ALD1 expression. MS-MS analysis supports structures related to hydroxyproline (Figures S5, compounds 5-7). While compound 3 has previously been reported as an FMO1-dependent product in pathogen-elicited *Arabidopsis* plants (Chen et al., 2018), none of these other molecules were detected in the native plant extracts analyzed in Figure 1C. It is possible that these putative NHP-related compounds only accumulate to detectable levels due to the high NHP concentration in *N. benthamiana* leaves expressing ALD1 and FMO1 and that proline hydroxylation is an artifact of FMO1 overexpression and not a native function of the enzyme.

### Engineering of *Arabidopsis* NHP pathway in *S. lycopersicum*

Transient expression of the NHP biosynthetic pathway in *N. benthamiana* not only demonstrates the minimum set of genes required for engineering NHP production, it also presents an opportunity to quickly determine if *biosynthetically* produced NHP could protect against pathogen infection to crop plants naïve to an initial infection. In other words, can SAR be established with the constitutive expression of just two biosynthetic genes, *ALD1* and *FMO1*? We chose to test this concept in tomato, given the importance of this plant as a vegetable crop and the fact that it is amenable to *Agrobacterium-*mediated transient gene expression.

Similar to the SAR assay developed to test the activity of synthetic NHP (Figure 2), we utilized an individual branch of a fully expanded tomato leaf. For each leaf, two top leaflets were infiltrated with *Agrobacterium* induction medium (Mock) or a 3×10^8^ cfu/ml suspension of *Agrobacteria* carrying *GFP* alone or *GFP* and the two *Arabidopsis* NHP pathway genes *ALD1* and *FMO1* (Figure 4A). At 48 hpi, we performed metabolite profiling using GC-MS to monitor Pip and NHP in *Agrobacteria-*infected top leaflets and uninfected bottom leaflets. We found that Pip and NHP levels were detected in both infected top leaflets (Figure 4B). Addition of exogenous Pip substrate was not required for production of NHP when *Arabidopsis* ALD1 and FMO1 are coexpressed in these leaflets (Figure S6). Significantly higher levels of NHP were present in the top leaflets expressing GFP+ALD1+FMO1 (GAF) compared to GFP alone (Figure 4B). Only low levels of Pip were detected in the uninfected bottom leaflets. Notably, increasing NHP production in the upper leaflets of tomato leaves provided protection in distal, bottom leaflets challenged with *Pst* (Figures 4C and 4D). Infection of bottom leaflets with *Pst* 48 h post-transient expression of *Arabidopsis* ALD1 and FMO1, and not GFP alone, in upper leaflets significantly reduced the titer of *Pst* in the bottom leaflets (Figure 4D). Transient expression of ALD1+FMO1 provided similar protection as ALD1+SARD4+FMO1 (Figure S7), indicating that overexpression of SARD4 is not necessary for initiating defense priming in tomato leaves under these conditions. Taken together, these data demonstrate that engineering a minimal set of enzymes for the *Arabidopsis* NHP pathway in *S. lycopersicum* not only enhances NHP production but also primes SAR in distal leaf tissue.

**Figure 4.**
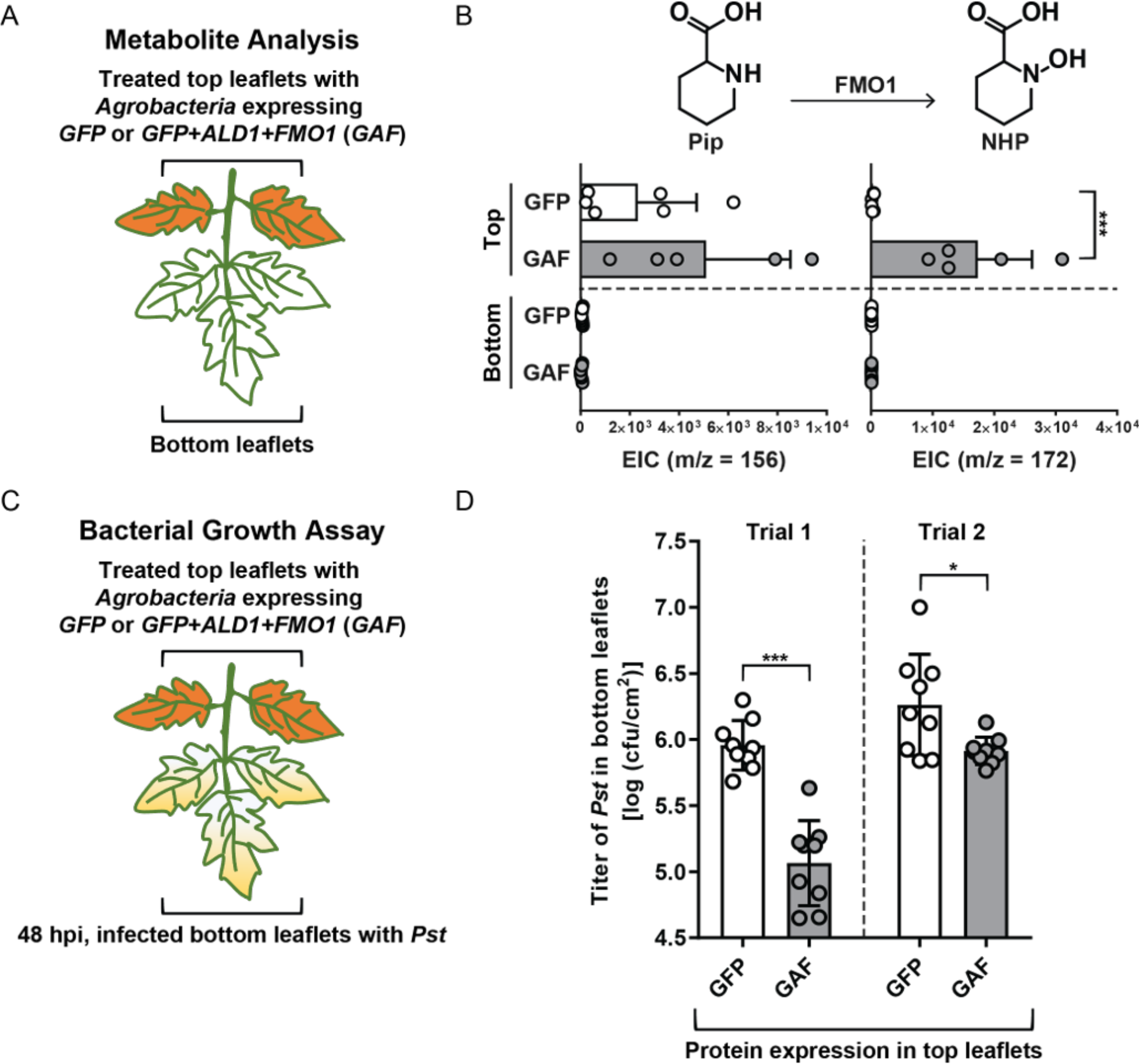
Engineering NHP pathway in *S. lycopersicum* cultivar VF36 leaves. (A) Diagram showing infection conditions for metabolite analysis. Two top leaflets of a tomato leaf were infiltrated with a suspension of *Agrobacteria* strains (final OD_600_ = 0.3) expressing *GFP* or *GFP* and two At NHP pathway genes (ALD1 + FMO1). At 48 hpi, the two top treated leaflets and three bottom untreated leaflets were harvested and analyzed by GC-MS. (B) Ion abundance of Pip (m/z = 156) and NHP (m/z = 172) in leaflets from (A). Bars of top leaves represent mean ± STD of 5 or 6 replicates. Bars of bottom leaves represent mean ± STD of 9 replicates. Levels reported as zero indicate no detection of metabolites. (C) Diagram showing infection conditions for bacterial growth assay. Two top leaflets of a tomato leaf were infiltrated with *Agrobacteria* strains as described in (A). At 48 hpi, the three bottom untreated leaflets were inoculated with *Pst* (1×10^5^ cfu/ml). The titer of *Pst* in the bottom leaflets was determined 4 days later. Bars represent mean ± STD of nine leaflets of three individual plants. Asterisks denote the significant differences between indicated samples using a one-tailed t-test (* *P* < 0.05; *** *P* < 0.001). The experiment was repeated twice with similar results.

## DISCUSSION

### Activity of FMO1 is conserved across phylogeny

The recent discovery that NHP is a bioactive metabolite that initiates SAR in *Arabidopsis* has provided new insight into the role of plant-derived chemical signals in propagating responses to biotic stress. Importantly, the presence of NHP in *Arabidopsis* has sparked questions about its biochemical evolution and function throughout the plant kingdom. NHP’s precursor, Pip, is known to enhance disease resistance in the Solanaceous plant *N. tabacum* (Vogel-Adghough et al., 2013), and has been reported to accumulate in other diseased angiosperms including *G. max* and *Oryza sativa* (Pálfi and Dézsi, 1968). In addition, Pip biosynthesis in the early vascular plant *Huperzia serrata* was found to proceed via homologs to the *Arabidopsis ALD1* and *SARD4* genes (Xu et al., 2018), suggesting that Pip biosynthesis, and perhaps NHP, is well-conserved across vascular plant phylogeny.

In this study, we provide bioinformatic and biochemical evidence that the production of NHP is conserved across vascular plants. We also provide biological evidence that pathogen-responsive accumulation of NHP occurs in *B. rapa*, a relative of *Arabidopsis* in the Brassicaceae and *N. benthamiana* and *S. lycopersicum*, two plants in the Solanaceae (Figures 1 and S2). Additionally, we show that transient expression of FMO1 homologs from diverse clades of dicots including Rosids (*A. thaliana, B. rapa, G. max*), Asterids (*S. lycopersicum*, *N. benthamiana*), and a monocot (*Z. mays*) in a heterologous host (*N. benthamiana*) catalyzes the conversion of Pip to NHP (Figures 1 and S3). Notably, the predicted patterns of NHP biosynthetic pathway conservation reflect that of two plant hormones with a well-established role in defense, SA and JA (Figure S1). Collectively, our data highlights the importance of NHP biosynthesis in plant species throughout the plant kingdom.

### Different plants may possess alternative processing mechanisms for NHP

We and others have reported NHP-Glc as an FMO1-dependent signal (Chen et al., 2018; Hartmann and Zeier, 2018) and have hypothesized that it may serve as a stable storage form of NHP, similar to the role that SA-glucose conjugates serve in defense priming (Dempsey et al., 2011). While we also observed high levels of NHP-Glc in *B. rapa* after *Pst* infection, we did not detect this signal in *N. benthamiana* or *S. lycopersicum*, even though free NHP accumulated in response to *Pst* (Figure S2). Interestingly, even in transient expression experiments in *N. benthamiana* and *S. lycopersicum*, where free NHP accumulates to levels 100-1000 fold higher than detected in *Pst*-elicited plants, no NHP-Glc was detected (Figures 3, S2 and S6). This is in contrast to findings in SA metabolism where SA-glucose conjugates have been detected in the Solanaceae and many other plant clades (Lee et al., 1995). These data suggest that there could be alternative NHP-processing mechanisms outside the Brassicaceae, and these processes may be important for signal activation, transport, or attenuation.

### Signal transport in *Arabidopsis* versus *S. lycopersicum*

We discovered that NHP-Glc accumulates in distal leaves of *Arabidopsis fmo1*, an NHP biosynthesis mutant, when local leaves are infiltrated with a solution of NHP (Chen et al., 2018). This suggests that NHP or an NHP derivative is transported systemically in *Arabidopsis*. By contrast, NHP and NHP-Glc were not detected in distal leaves of *S. lycopersicum* under our SAR assay conditions. These findings suggest that NHP’s stability, derivatization and/or transport may be distinct within Solanaceous plants. Despite our inability to detect NHP accumulation in distal tomato leaflets, local infiltration of NHP provided enhanced protection against bacterial infection in distal leaflets (Figure 2), which implies that either an unidentified NHP derivative is being transported or that NHP is inducing production and/or transport of a different mobile signal to prime defense. Recent evidence in *Arabidopsis* indicates that, while some Pip appears to be transported, Pip confers SAR by increasing nitric oxide and reactive oxygen species, which can trigger AzA/G3P-associated SAR (Wang et al., 2018). A similar mechanism could be occurring in response to Pip/NHP in *S. lycopersicum*, which would not necessitate the transport of Pip/NHP to confer resistance.

### Engineering NHP biosynthesis for disease protection

Several aspects of plant defense, including pathogen detection, immune signaling, and antimicrobial production have been the targets of engineering in an effort to enhance native immunity in crops. For example, since the discovery that the *Arabidopsis* elongation factor Tu receptor (EFR) could function in Solanaceous plants (Lacombe et al., 2010), many immune receptor genes have been transferred between plant families to expand native pathogen detection (Rodriguez-Moreno et al., 2017). An alternative strategy for engineering disease resistance has focused on introducing biosynthetic genes that produce small-molecule phytoalexins with direct antimicrobial activities (Großkinsky et al., 2012). This approach has been successful, especially with the well-studied phytoalexin resveratrol, which has been engineered into numerous plants to enhance pathogen resistance (Delaunois et al., 2008). Other small molecule-based approaches have utilized direct chemical priming, where application of chemical inducers enhances native disease mechanisms in a manner akin to SAR. This approach allows bypassing of species-specific pathogen detection, instead relying on external induction of a plant’s native defense response. Such a strategy could enable protection against the emergence of a pathogen before contact with a crop plant and/or enhanced resistance against adapted pathogens that actively suppress the plant host defense responses directly downstream of detection. Several synthetic chemical inducers of this type have met success in commercial agriculture and are being utilized in combination with antimicrobials in the field (Conrath et al., 2015). It is notable that these commercial inducers (such as benzothiadiazole (Görlach et al., 1996)) are entirely synthetic and cannot be made using engineered biosynthesis. In contrast, the production of natural inducers of SAR, including NHP, could be engineered to elicit potent responses to pathogen infections. External control of SAR induction allows for activation of native defense responses before the onset of infection. Each of these approaches suffers from different drawbacks. For example, several are vulnerable to the emergence of pathogen resistance. However, deploy a combination of multiple strategies may be the most effective way to achieve durable disease resistance.

The discovery that diverse plant species have the ability to produce NHP and that its exogenous application can enhance protection against bacterial infection in *S. lycopersicum* and *C. annuum* hold promise for future efforts to engineer disease resistance in important crop plants. While *S. lycopersicum* is capable of naturally producing NHP, we have shown that exogenous application can bypass the need for an initial infection to trigger an SAR defense response (Figure 2). By transiently expressing *Arabidopsis* ALD1 and FMO1 in *N. benthamiana*, NHP production can be increased 100-1000 fold over accumulation in native plants (Figures 3 and S2), so it follows that by engineering plants to overproduce NHP, disease resistance could be enhanced. As a proof-of-concept, we have shown that the NHP pathway can be engineered under a high expression constitutive promoter, and other stable engineering approaches could utilize external chemical inducers or natural plant regulatory systems to minimize growth/yield tradeoffs that may come with NHP production (Ning et al., 2017).

There are likely to be challenges associated with using the NHP pathway to engineer resistance, but with these challenges come opportunities to probe fundamental questions in SAR, including how small molecules interact to establish functional SAR, how signals are transported, perceived, and attenuated, and how SAR mechanisms have been conserved and diversified throughout plant evolution. Collectively, our data provide insight into the function of NHP across vascular plants and highlight the importance of further study into SAR biology and chemical defenses in agriculturally relevant crops.

## ACKNOWLEDGEMENTS

We thank George Lomonossoff (John Innes Centre) for providing pEAQ plasmid. This work was supported by an HHMI and Simons Foundation Grant 55108565 (to E.S.S.), National Science Foundation Graduate Research Fellowship DGE-1656518 (to E.C.H.), National Science Foundation IOS-1555957 and Binational Science Foundation Grant 2011069 (to M.B.M.) and Ministry of Science and Technology of Taiwan-105-2917-I-564-414 093 (to Y.C.C.).

## AUTHOR CONTRIBUTIONS

E.C.H., Y.C.C., E.S. and M.B.M. contributed to the study design; E.C.H. and Y.C.C. performed research; E.C.H., Y.C.C., E.S. and M.B.M. analyzed data; and E.C.H., Y.C.C., E.S. and M.B.M. wrote the manuscript.

## DECLARATION OF INTERESTS

Nothing to declare.

## EXPERIMENTAL MODEL

### Plant materials and growth conditions

For seedling hydroponics experiments, 15 ± 1 *Arabidopsis thaliana* ecotype Col-0 seeds, 2 rapid-cycling *Brassica rapa* seeds, 2 *S. lycopersicum* Heinz 1706 seeds, 10 ± 1 *N. benthamiana* Nb-1 seeds, 1 *Glycine max* Pella 86 seed, and 1 *Zea mays* B73 seed, respectively, were placed into 3 ml of sterile 1x Murashige-Skoog (MS) medium with vitamins (PhytoTechnology Laboratories) (pH 5.7) + 5 g/l sucrose in the wells of a 6-well microtiter plate. Plates were kept in the dark at room temperature for 48 h and then transferred to a growth chamber at 50% humidity, 22°C, and 100 μmol/m^2^/s photon flux under a 16-h light/ 8-h dark cycle. After 1 week of growth, spent medium was replaced with 3 ml of fresh MS medium + 5 g/l sucrose. *B. rapa*, *S. lycopersicum*, *G. max*, *and Z. mays* seedlings were elicited at 1 week. *A. thaliana* and *N. benthamiana* seedlings were elicited at 2 weeks. For adult plant experiments, tomato (*Solanum lycopersicum* cultivar VF36), pepper (*Capsicum annuum* cultivar Early Calwonder) and *Nicotiana benthamiana* plants were grown in a greenhouse (16-h light / 8-h dark, 25–28°C) and used at 4-6 weeks of age.

### Bacterial strains and growth conditions

*Escherichia coli* strains DH5 alpha and 10-beta, *Pseudomonas syringae* pathovar *tomato* strain DC3000 (*Pst*), *Xanthomonas euvesicatoria* strain *Xe* 85-10 and *Agrobacterium tumefaciens* strains GV3101 and C58C1 pCH32 were used in this study. *E. coli* strains were grown in Lysogeny broth (LB) agar containing appropriate antibiotics at 37°C. *Pseudomonas* and *Xanthomonas* strains were grown at 28°C on nutrient yeast glycerol agar (NYGA) medium containing rifampicin 100 μg/ml. *Agrobacteria* strains were grown at 28°C on LB agar containing rifampicin 100 μg/ml, tetracycline 5 μg/ml and kanamycin 50 μg/ml for C58C1 pCH32 and gentamycin 100 μg/ml and kanamycin 50 μg/ml for GV3101.

## METHOD DETAILS

### Elicitation methods for seedling hydroponics experiments

*Pst* was grown on LB agar plates at 30°C overnight. A single colony was picked and grown to mid-log phase in liquid LB at 30°C. Cells were centrifuged, washed three times with MS + 5 g/l sucrose, and resuspended in MS + 5 g/l sucrose to an OD_600_ of 0.2. 20 μl of sterile 10 mM MgCl_2_ was added to wells of mock-elicited samples and 20 μl of the *Pst* suspension was added to each treatment well. Plants were elicited 48 h prior to sample harvest.

### Plant tissue extraction for metabolite profiling

All plant tissue was flash frozen, lyophilized to dryness, and homogenized at 25 Hz for 2 min using a ball mill (Retsch MM 400). Samples were resuspended in 20 μl 80:20 MeOH:H_2_O per mg of dry tissue and incubated at 4°C for 10 min. The liquid fraction of each sample was split, and one half was passed through a 0.45 μm PTFE filter and directly analyzed via liquid chromatography-mass spectrometry (LC-MS). The other half was dried under an N_2_ stream and further processed for gas chromatography-mass spectrometry (GC-MS) analysis. Dried samples were resuspended in 250 μl of a 10% v/v N-Methyl-N-(trimethylsilyl)trifluoroacetamide (MSTFA) with 1% trimethylchlorosilane (Sigma-Aldrich) solution in hexanes, briefly vortexed, and incubated at 70°C for 30 min. Samples were then allowed to cool to room temperature, filtered through 0.45 μm PTFE filters, and analyzed via GC-MS.

### Cloning of plant *FMO1* homologs and *A. thaliana* NHP biosynthetic genes

*FMO1* homologs from *S. lycopersicum* cultivar VF36 and *N. benthamiana* were PCR amplified using cDNA isolated from plant leaves and gene specific primers (Table S1) and then cloned into the pCR8/GW/TOPO vector (Life Technologies, Carlsbad, CA). cDNA sequences were confirmed by DNA sequence analysis and then subcloned into pEAQ-HT-DEST3 (Peyret and Lomonossoff, 2013) to create C-terminal 6xHis-tagged fusion proteins. Plasmids were then transformed into *E. coli* DH5 alpha and *A. tumefaciens* strain C58C1 pCH32 by heat shock transformation.

*FMO1* cDNA homologs from *B. rapa* (NCBI: XP_009149401.1; BraA06g014860), *G. max* (NCBI: XP_003541317.1; Glyma13g17340), and *Z. mays* (NCBI: XP_008660479.1; AC191071.3_FGP001) were codon optimized for expression in *N. benthamiana* using Integrated DNA Technologies’ codon optimization tool (https://www.idtdna.com/codonopt) (Supplementary information), which eliminates codons with a usage rate <10%, assigns codons with a frequency matching that of the target organism, and further optimizes to streamline synthesis of the gene. The optimized genes were synthesized as gBlock fragments (Integrated DNA Technologies), cloned into pEAQ-HT (Peyret and Lomonossoff, 2013) under the control of the CaMV 35S promoter, and transformed into *E. coli* 10-beta and *A. tumefaciens* GV3101 by heat shock transformation.

*A. thaliana ALD1*, *SARD4*, and *FMO1* cDNAs were amplified by PCR using *A. thaliana* cDNA and gene-specific primers (Table S1). These products were cloned into pEAQ-HT (Peyret and Lomonossoff, 2013) under control of the CaMV 35S promoter and transformed into *E. coli* 10-beta, *A. tumefaciens* GV3101, and *A. tumefaciens* C58C1 pCH32 by heat shock transformation.

### Transient expression in *N. benthamiana*

*A. tumefaciens* strains engineered with NHP biosynthetic gene constructs were grown for 48 h on LB agar plates with appropriate antibiotics. Cells were scraped from the surface of the plate using an inoculation loop, washed three times, resuspended in *Agrobacterium* induction medium (10 mM MES buffer, 10 mM MgCl_2_, 150 μM acetosyringone, pH 5.7), and incubated at room temperature with shaking for 2 h. For transient expression of plant *FMO1* homologs, cells were diluted to a final OD_600_ of 0.3 in induction medium + 1 mM Pip. For transient expression of *A. thaliana* NHP pathway genes, cells were combined in equal proportions with a *GFP* expressing strain for a final OD600 of 0.4 in induction medium. These solutions were infiltrated into the underside of *N. benthamiana* leaves using a needleless 1 ml syringe. Plants were returned to the growth shelf for 24 h prior to tissue harvest and processing.

### LC-MS metabolite profiling

NHP-Glc was measured using a previously-documented reverse-phase chromatography method on an Agilent 1260 HPLC coupled to an Agilent 6520 Q-TOF ESI mass spectrometer (Chen et al., 2018).

### GC-MS metabolite profiling

Plant extracts derivatized with MSTFA were analyzed on an Agilent 7820A gas chromatograph coupled to an Agilent 5977B mass spectrometer. Samples were flown through a fused silica column (Agilent VF-5ht, 30 m × 0.25 mm × 0.1 μm) using the following oven temperature gradient: 70°C for 2 min, ramp to 150°C at 10°C/min, and final hold at 320°C for 2 min. To detect Pip (2-trimethylsilyl (2-TMS) derivative) and NHP (2-TMS derivative), the instrument was run in both scan and selected ion monitor (SIM) modes where the selected ions were m/z 156, 230, and 273 for Pip and m/z 172, 246, and 274 for NHP. Reported values for ion abundance are peak integration values of extracted ion chromatograms for Pip (m/z 156) and NHP (m/z 172) detected in SIM mode. Fragmentation patterns and retention times for *in planta* Pip and NHP were verified with authentic standards of Pip (Oakwood Chemical) and NHP (Chen et al., 2018).

### Transient expression of *Arabidopsis* NHP biosynthesis genes in tomato

*A. tumefaciens* strains C58C1 pCH32 carrying *A. thaliana* NHP biosynthetic gene constructs (*ALD1, FMO1* and/or *SARD4*) or *an Arabidopsis FMO1* inactive FAD binding mutant *(At FMO1mutant, G17A/G19A) or S. lycopersicum FMO1 (Sl FMO1)* were grown on LB agar plates with appropriate antibiotics. Cells were scraped from the surface of the plate, washed 1-2 times, resuspended in *Agrobacterium* induction medium (10 mM MES buffer, 10 mM MgCl_2_, 150 μM acetosyringone, pH 5.7), and incubated at room temperature for 2 h. Cells of a single strain were inoculated at a final OD_600_ of 0.1 in induction medium. Cells of different strains were combined for a final OD_600_ of 0.3 in induction medium. Two leaflets of 3^rd^ or 4^th^ compound leaves of 4 to 5-week old *S. lycopersicum* VF36 plants were infiltrated with each suspension (one suspension per plant). Plants were placed under light (16-h light/ 8-h dark or continuous light) for 24 h or 48 h prior to tissue harvest or bacterial growth assays. For bacterial growth assays, three terminal leaflets on the same branch were inoculated with a 1×10^5^ cfu/ml suspension of *Pst*. Plants were then incubated in a greenhouse (16-h light/ 8-h dark, 25–28°C). The titer of *Pst* in the terminal leaflets was quantified 2 or 4 days post-infiltration (dpi) by homogenizing leaf discs in 1 ml 10 mM MgCl_2_, plating appropriate dilutions on NYGA media with rifampicin (100 μg/ml), incubating plates at 28°C for 2 days, and then counting bacterial colonies. Three biological repeats were performed per treatment in two independent experiments.

### SAR assay in tomato

Two top leaflets (closest to main stem) of 3^rd^ or 4^th^ compound leaves of 4 to 5-week old *S. lycopersicum* VF36 plants were infiltrated with 10 mM MgCl_2_ (Mock) or 10 mM MgCl_2_ containing 1 mM NHP. After chemical incubation for 24 h, three bottom leaflets (distal to main stem) on the same branch were inoculated with a 1×10^5^ cfu/ml suspension of *Pst*. Plants were incubated in greenhouse (16-h light/ 8-h dark, 25–28°C). The titer of *Pst* in the bottom leaflets was quantified at 4 dpi by homogenizing leaves discs in 1 ml of 10 mM MgCl_2_, plating appropriate dilutions on NYGA media with rifampicin (100 μg/ml), incubating plates at 28°C for 2 days, and then counting bacterial colonies. Symptoms of infected leaves were photographed at 4 dpi. Three biological repeats were performed per treatment in two independent experiments.

### SAR assay in pepper

Two bottom leaves of 4 to 5-week old pepper plants were infiltrated with 10 mM MgCl_2_ or 10 mM MgCl_2_ containing 2 mM NHP. After chemical incubation for 24 h, one upper leaf was inoculated with a 1×10^4^ cfu/ml suspension of *Xe* 85-10. Plants were incubated in a greenhouse (16-h light/ 8-h dark, 25–28°C). The titer of *Xe* 85-10 in the infected leaves was quantified 5 and 10 dpi by homogenizing leaves discs in 1 ml 10 mM MgCl_2_, plating appropriate dilutions on NYGA media with rifampicin (100 μg/ml), incubating plates at 28°C for 2 days, and then counting bacterial colonies. Symptoms of infected leaves were photographed at 10 dpi. Four biological repeats were performed per treatment in two independent experiments.

## SUPPLEMENTARY FIGURES

**Figure S1.**
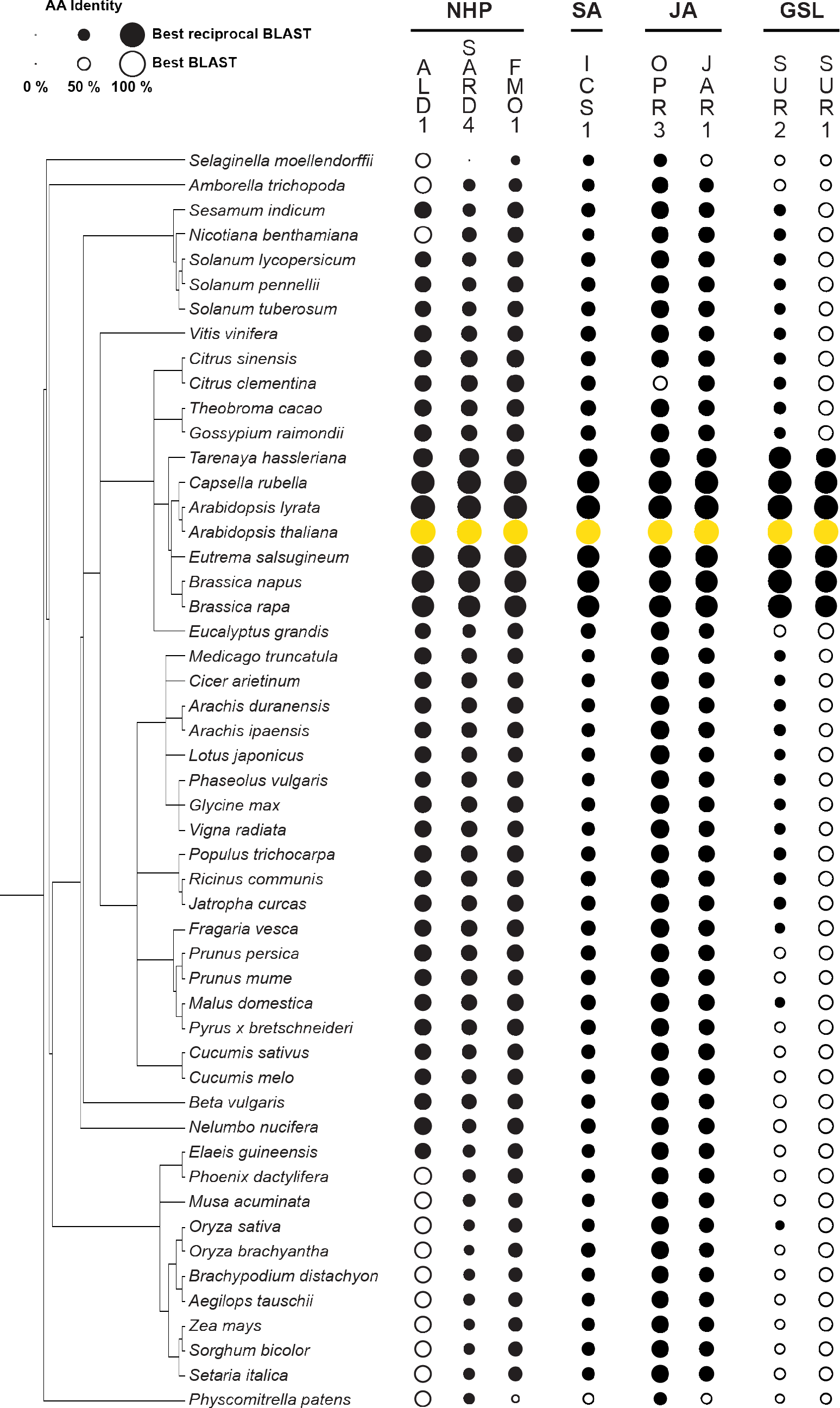
BLAST analysis of plant defense metabolites (Related to Figure 1) Phylogenetic tree of sequenced plant genomes created using the PhyloT tree generator (http://phylot.biobyte.de/index.html). Circles are scaled by the percent amino acid identity of the best BLAST (blastp) hit between the *Arabidopsis thaliana* biosynthetic proteins (yellow) and the homologs of the respective plant proteome (black). Solid circles signify that the homolog returned the respective *A. thaliana* protein in a reverse-BLAST into the *A. thaliana* proteome (best reciprocal BLAST). Biosynthetic pathways analyzed are NHP, salicylic acid (SA), jasmonic acid (JA), and glucosinolate (GSL). Specifically, we analyzed isochorismate synthase 1 (ICS1; (Wildermuth et al., 2001)) for SA, OPDA reductase (OPR3; (Stintzi and Browse, 2000)) and jasmonate resistance 1 (JAR1; (Staswick and Tiryaki, 2004)) for JA, and S-alkyl thiohydroximate lyase (SUR1; (Mikkelsen et al., 2004)) and cytochrome P450 monooxygenase CYP83B1 (SUR2; (Bak et al., 2001)) for GSL. For the SA biosynthetic enzyme ICS1, reverse BLASTs returning *A. thaliana* ICS1 and isochorismate synthase 2 (ICS2; a functionally redundant enzyme (Macaulay et al., 2017)) were considered best reciprocal BLASTs. The analysis predicts that SA and JA pathways are highly conserved throughout the plant kingdom, while the GSL pathway is only conserved within the Brassicaceae.

**Figure S2.**
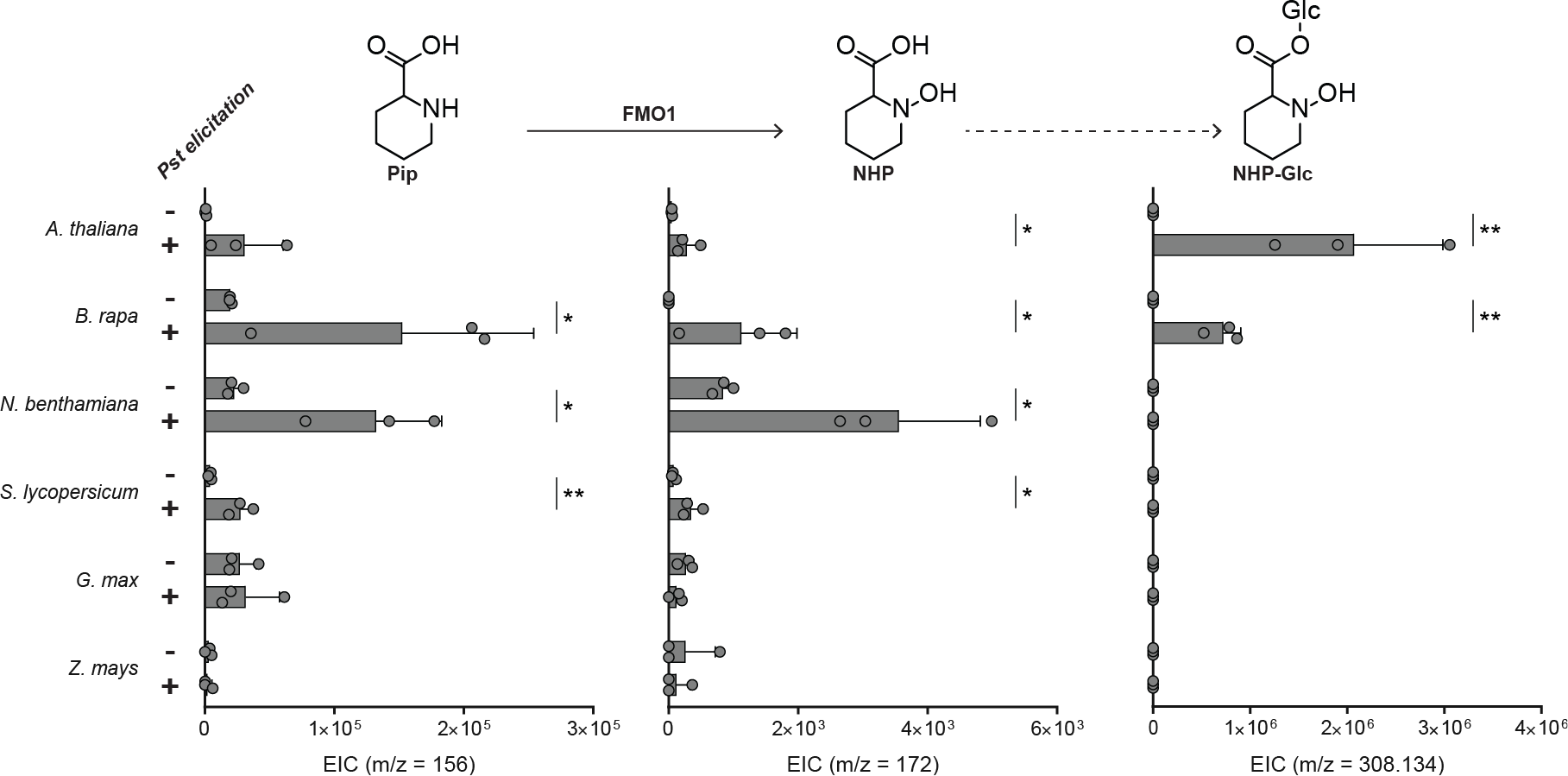
Production of Pip, NHP, and NHP-Glc in seedlings in response to *Pst* elicitation (Related to Figure 1) Ion abundances of Pip (m/z = 156), NHP (m/z = 172), and NHP-Glc (m/z = 308.134) detected by GC-MS (Pip, NHP) and LC-MS (NHP-Glc) in extracts isolated from *A. thaliana*, *B. rapa*, *N. benthamiana*, *S. lycopersicum*, *G. max*, and *Z. mays* seedlings elicited with 10 mM MgCl_2_ (− *Pst* elicitation) or *Pseudomonas syringae* pathovar *tomato* strain DC3000 (+ *Pst* elicitation). Levels represent the mean +/− STD of three biological replicates. Asterisks denote levels that significantly increased with *Pst* elicitation (one tailed t-test; * *P* < 0.05, ** *P* < 0.01). Bars represent the mean ± STD of three biological replicates. Levels reported as zero indicate no detection of metabolites.

**Figure S3.**
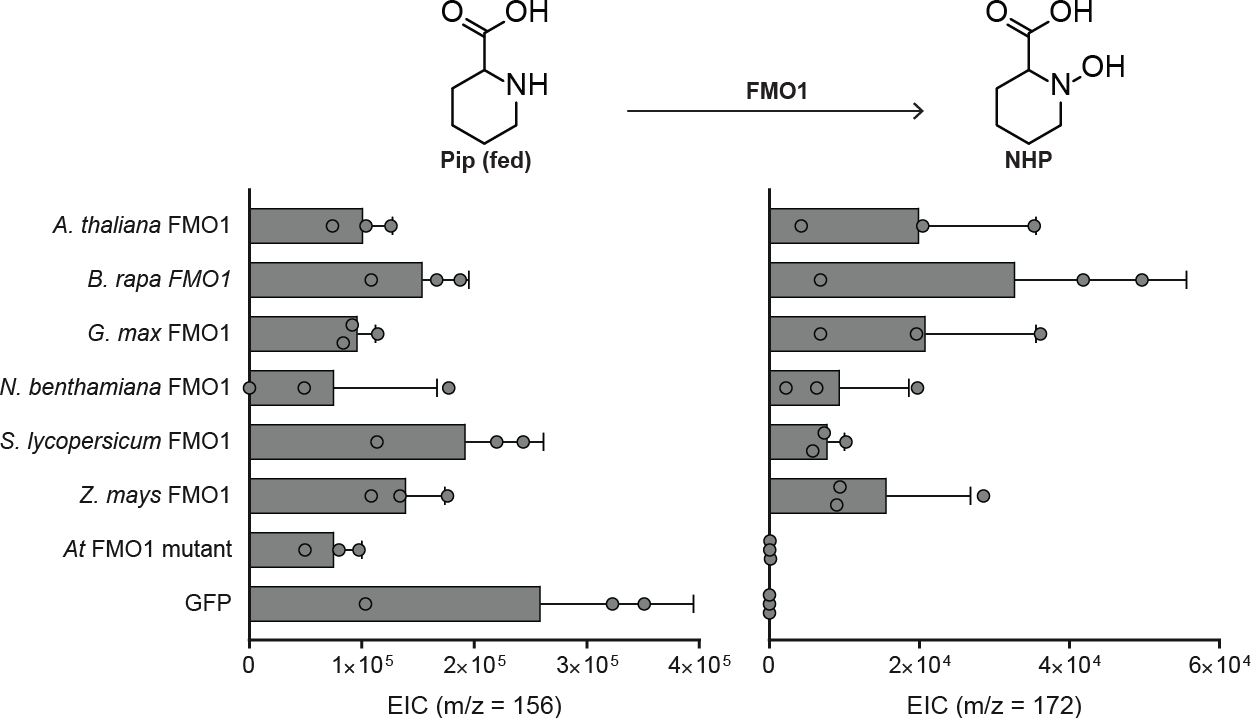
Production of NHP catalyzed by FMO1 homologs transiently expressed in *N. benthamiana* leaves (Related to Figure 1) Ion abundance of Pip (m/z = 156) and NHP (m/z =172) in *Agrobacterium* infected *N. benthamiana* leaves expressing *A. thaliana* FMO1, the most similar FMO1 homolog from *B. rapa* (mustard), *G. max* (soybean), *S. lycopersicum* (tomato), *Z. mays* (corn), an *A. thaliana* FMO1 mutant (G17AG19A; catalytically inactive) or GFP. 1 mM Pip was co-infiltrated with all *Agrobacteria* strains. Bars represent the mean +/− STD of three biological replicates harvested 24 hpi. Levels reported as zero indicate no detection of metabolites.

**Figure S4.**
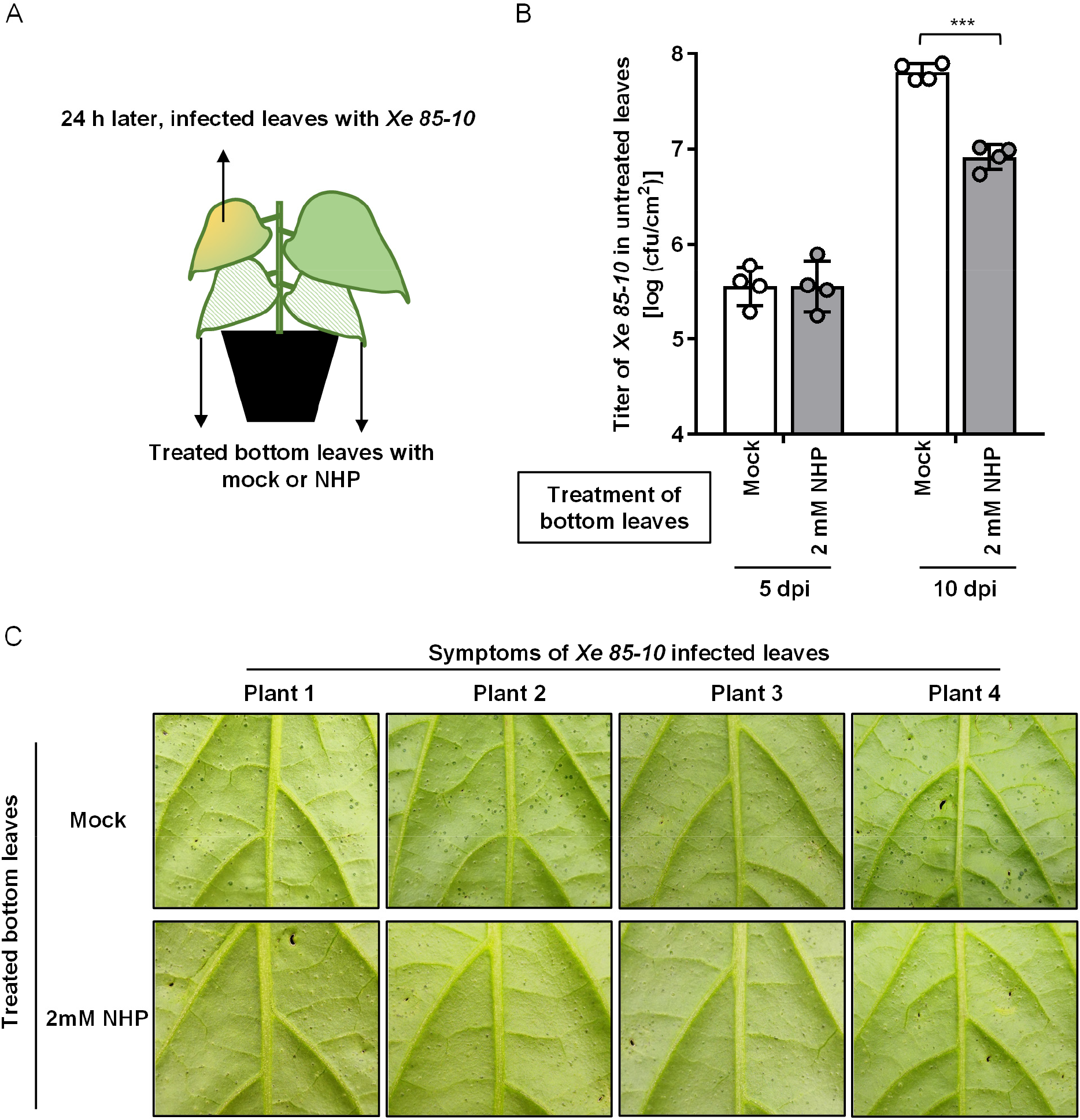
Local treatment of NHP induces systemic defense in pepper, *Capsicum annuum* cultivar Early Calwonder plants (Related to Figure 2) (A) The diagram showing the representative leaves used for SAR experiments. Two bottom leaves were infiltrated with 10 mM MgCl_2_ (Mock) or 10 mM MgCl_2_ containing 2 mM NHP. After chemical incubation for 24 h, one top leaf was inoculated with a 1×10^4^ cfu/ml suspension of *Xanthomonas euvesicatoria* strain 85-10 (*Xe 85-10*). (B) Titer of *Xe 85-10* in challenged leaves of pepper plant at 5 and 10 dpi. Bars represent the mean +/− STD of 4 biological replicates. Asterisks denote the significant differences between indicated samples using a one-tailed t-test *** P< 0.001). The experiment was repeated twice with similar results. (C) Symptoms of pepper leaves infected with *Xe 85-10*. Back side of leaves were photographed at 10 dpi and enlarged sections are shown to see pustule disease lesions and healthy tissue.

**Figure S5.**
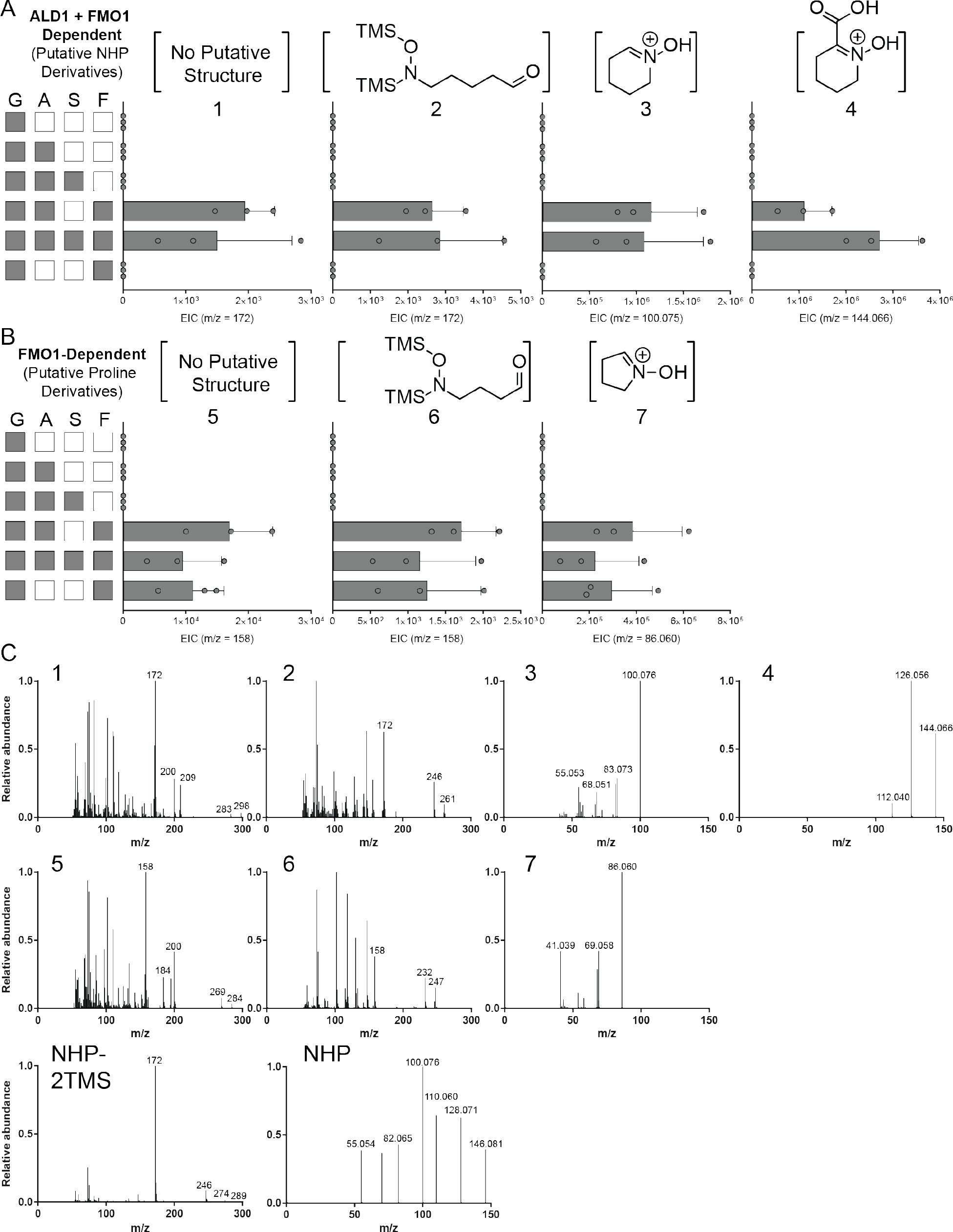
Levels of unknown compounds during transient expression of *Arabidopsis* NHP pathway genes in *N. benthamiana* (Related to Figure 3) Ion abundances of (A) unknown ALD1 and FMO1-dependent compounds and (B) unknown FMO1-dependent compounds in *N. benthamiana* leaves expressing combinations of *Arabidopsis ALD1* (abbreviation A), *SARD4* (S) *FMO1* (F) and *GFP* (G) genes. Each row represents a different enzyme set where a grey box indicates that an *Agrobacterium* strain containing that gene was included in the infiltration. For each biosynthetic gene, the *Agrobacterium* strain was infiltrated at an OD_600_ of 0.1 and a strain harboring *GFP* was included to normalize the total OD_600_ of infiltration at 0.4. Bars represent the mean +/− STD of three biological replicates harvested at 24 hpi. Levels reported as zero indicate no detection of metabolites. (C) MS/MS analysis of unknown compounds in (A) and (B). Experimental parameters for compounds are as follows: 1 (GC-MS, retention time (rt) = 9.41 min); 2 (GC-MS, rt = 6.55 min); 3 (RP LC-MS, rt = 0.9 min, collision induced dissociation (CID) = 40V); 4 (RP LC-MS, rt = 1.84 min, CID = 10V); 5 (GC-MS, rt = 8.25 min); 6 (GC-MS, rt = 5.83 min); 7 (RP LCMS; rt = 0.9 min; CID = 40V); NHP-2TMS (GC-MS, rt = 9.2 min); NHP (CID = 10V, included for reference (Chen et al., 2018)).

**Figure S6.**
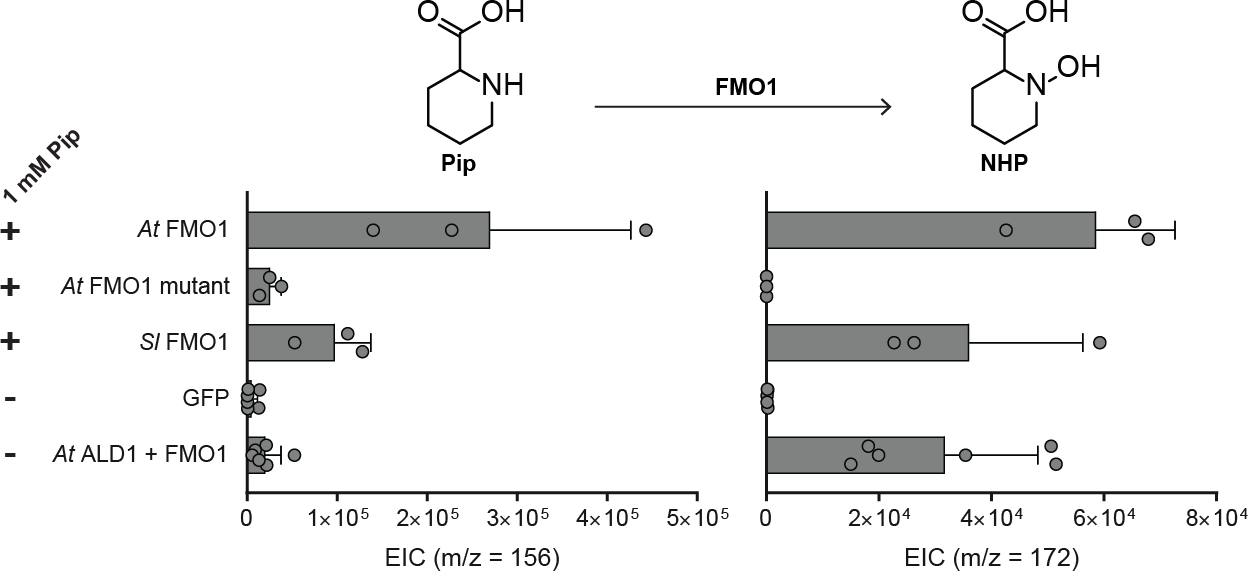
Production of NHP during transient expression in *S. lycopersicum* cultivar VF36 leaves (Related to Figure 4) Ion abundance of Pip (m/z = 156) and NHP (m/z = 172) in *S. lycopersicum* leaflets expressing *Arabidopsis* FMO1 (*At* FMO1), an *Arabidopsis* FMO1 inactive FAD binding mutant (*At* FMO1 mutant, G17A/G19A), *S. lycopersicum* FMO1 (*Sl* FMO1), GFP, or the minimal *Arabidopsis* NHP pathway (ALD1/FMO1/GFP). *A. tumefaciens* strains were infiltrated with (+) or without (−) 1 mM Pip. *S. lycopersicum* leaves were harvested at 72 hpi for GC-MS analysis. Bars represent the mean ± STD of three or six biological replicates. Levels reported as zero indicate no detection of metabolites.

**Figure S7.**
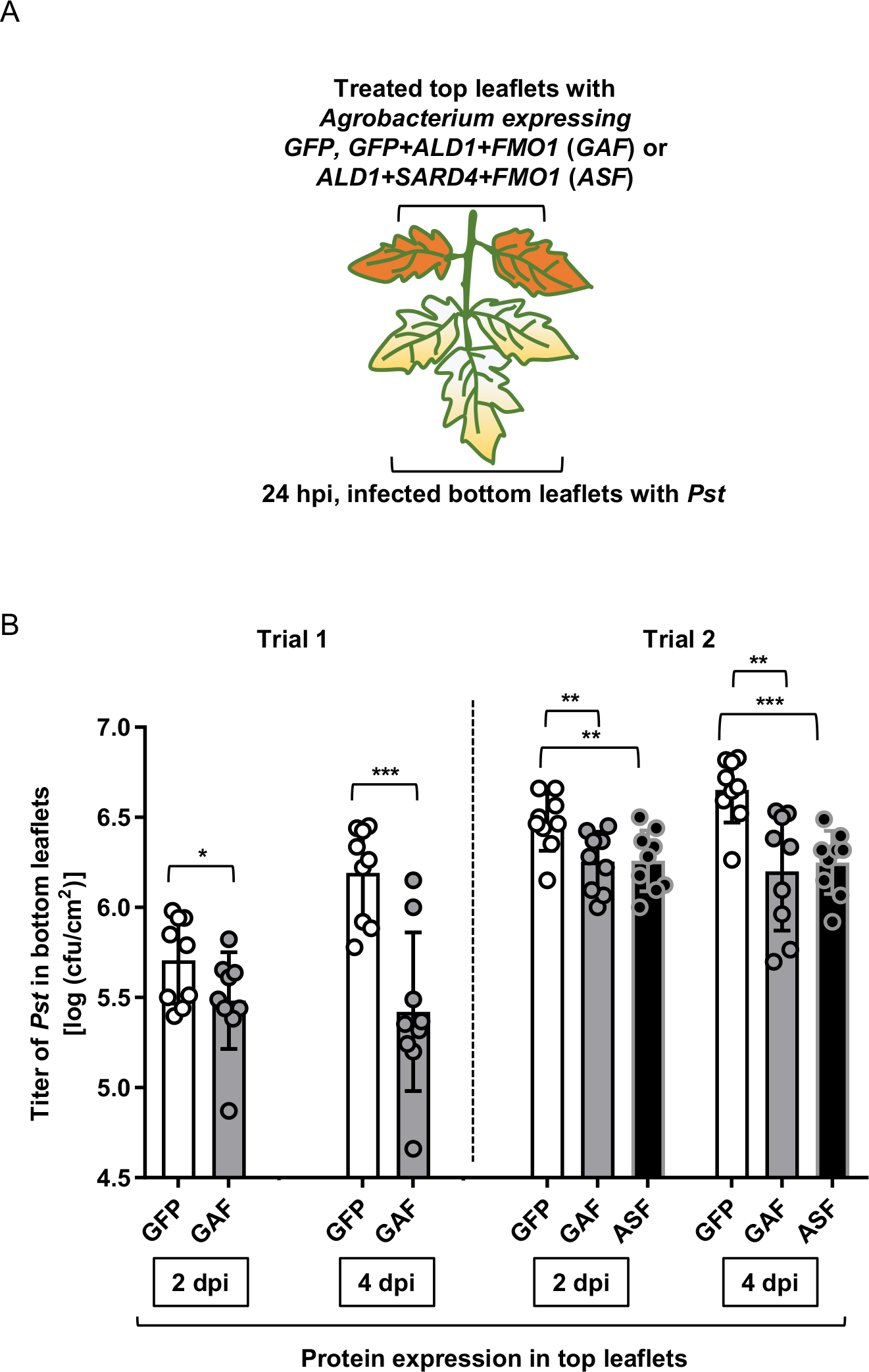
Engineering NHP pathway in *S. lycopersicum* cultivar VF36 leaves (Related to Figure 4) (A) Diagram showing infection conditions for bacterial growth assay.). Two top leaflets of a tomato leaf were infiltrated with a suspension (final OD_600_ = 0.3) of *Agrobacteria* expressing GFP or enzymes from the *Arabidopsis* NHP pathway [GFP + ALD1 + FMO1 (GAF) or ALD1 + SARD4 + FMO1 (ASF)]. At 24 hpi, three bottom untreated leaflets were inoculated with a 1×10^5^ cfu/ml suspension of *Pst.* (B)The titer of *Pst* in the infected bottom leaflets was determined at 2 and 4 days post *Pst* infection (dpi). Bars represent mean ± STD of nine leaflets of three individual plants. Asterisks denote the significant differences between indicated samples using a one-tailed t-test (* *P* < 0.05; ** *P* < 0.01; *** *P* < 0.001).

### Supplemental information

#### Sequences of codon-optimized FMO1 genes

*B. rapa FMO1* (optimized from XP_009149401.1):

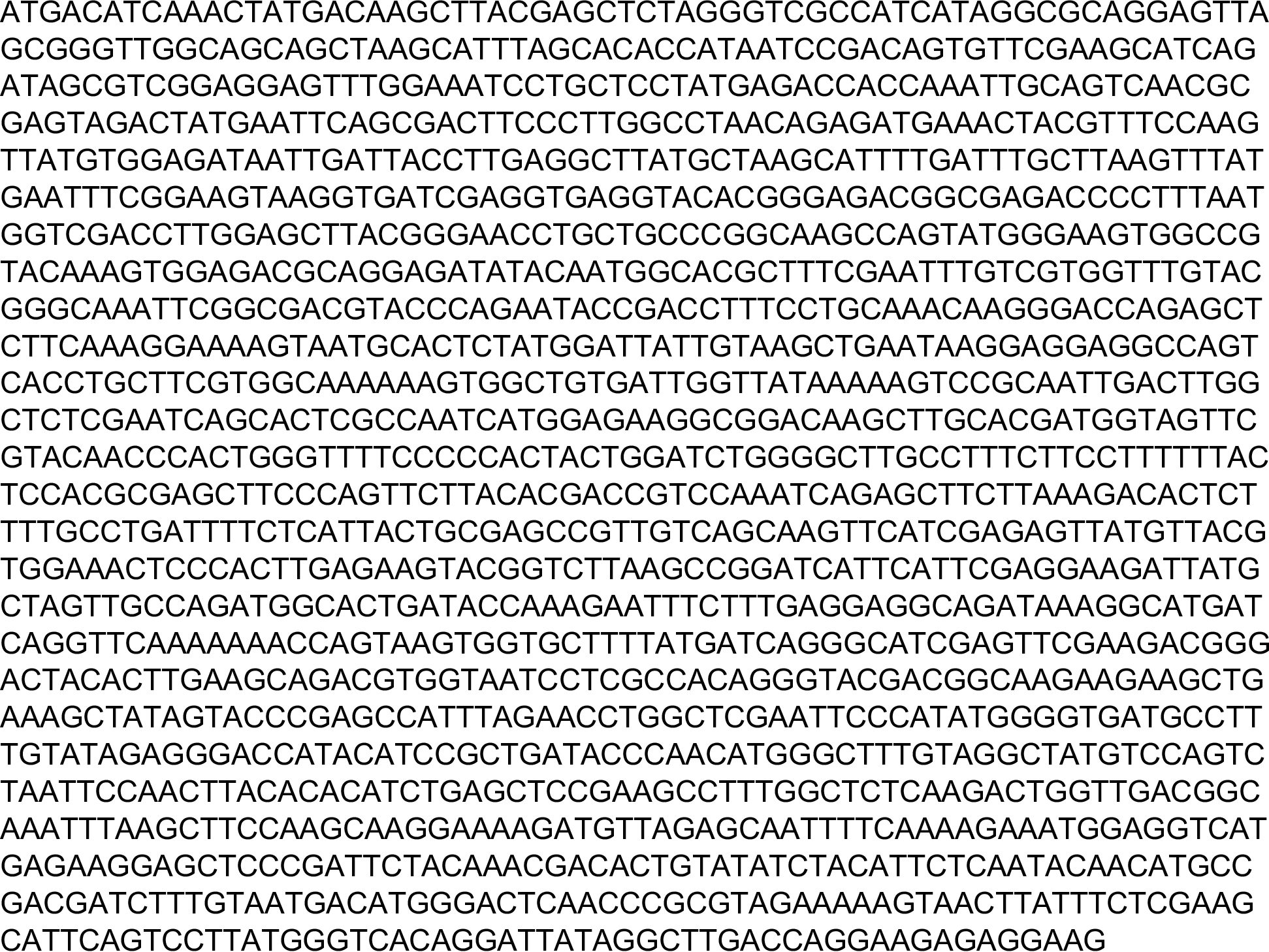

*G. max FMO1* (optimized from XP_003541317.1):

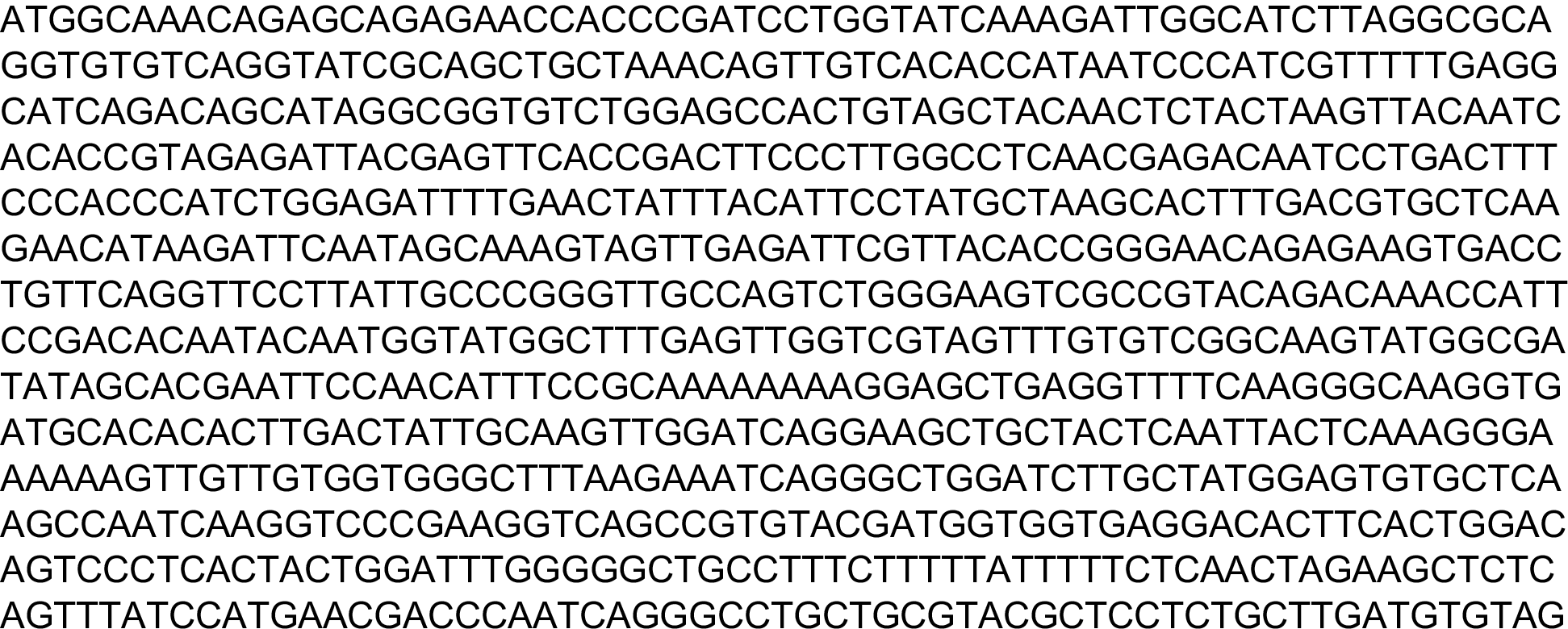

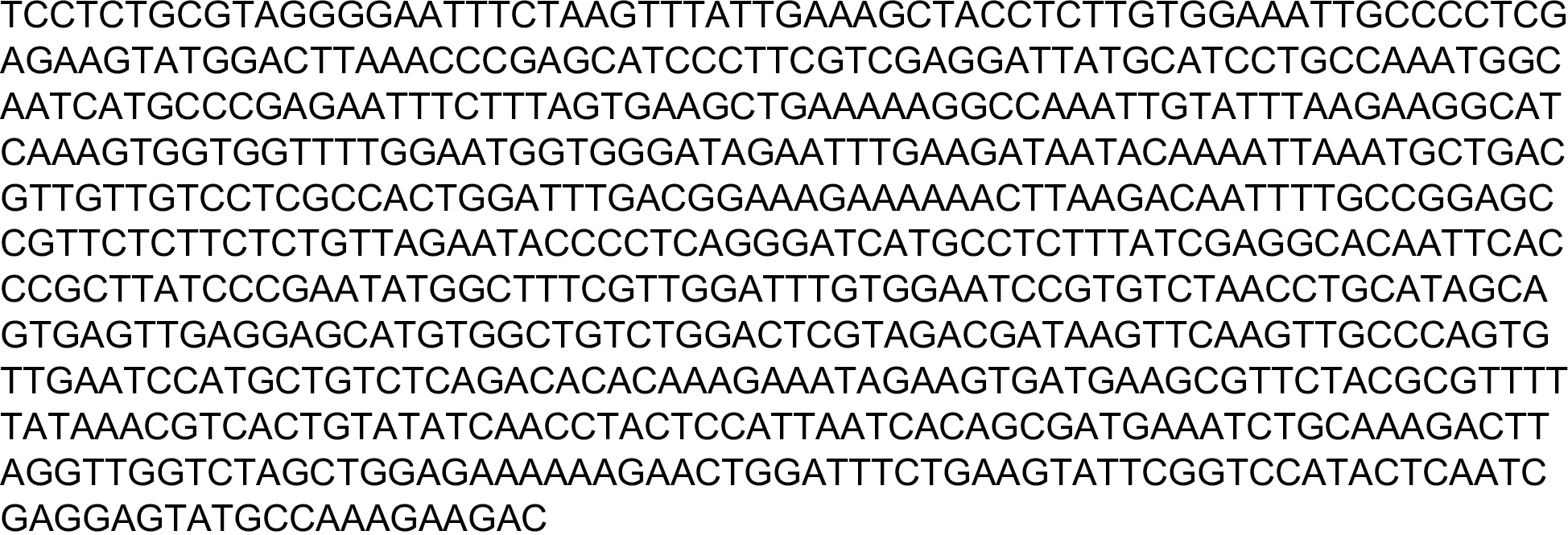

*Z. mays FMO1* (optimized from XP_008660479.1):

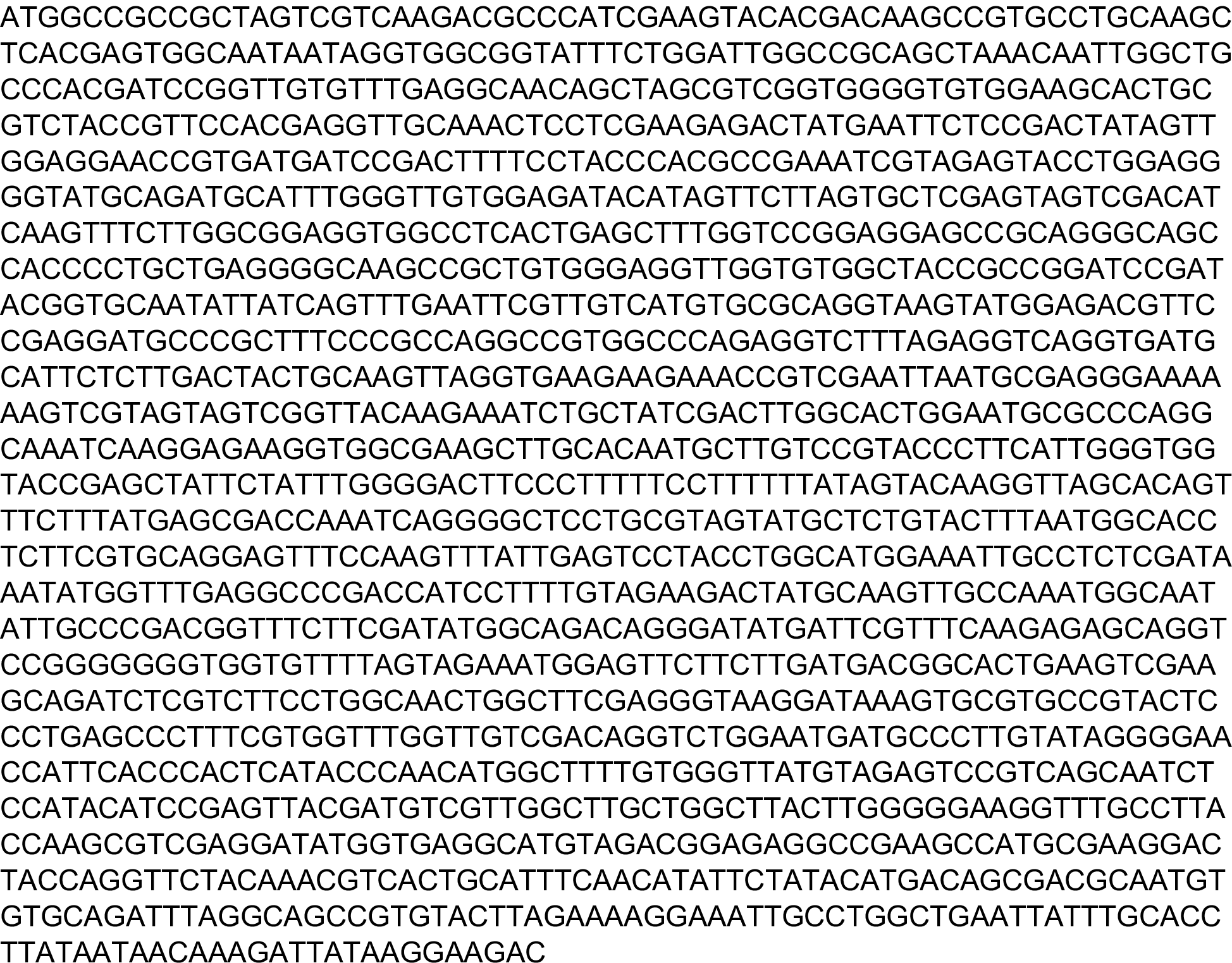

**Supplementary Table 1.**
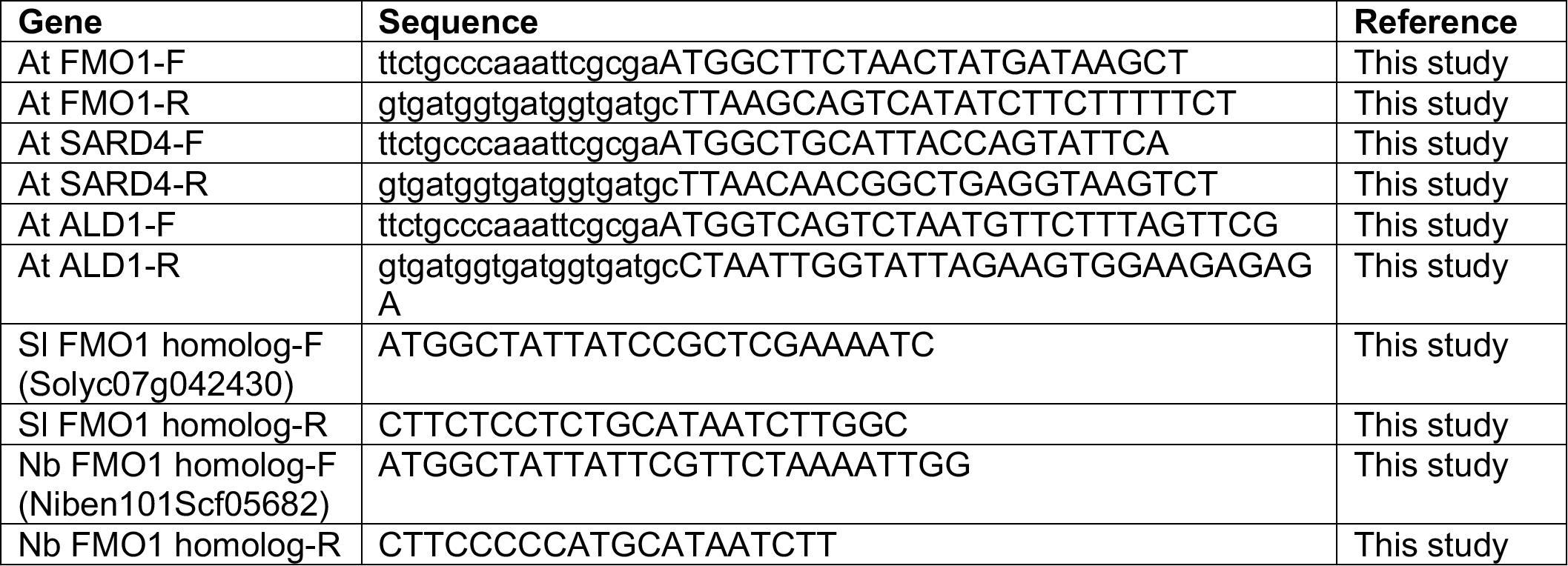
Primers used in this study

**Supplementary Table 2.** Percent amino acid identity of best BLAST hits to *A. thaliana* proteins. Percent amino acid identities of best BLAST hits shown in Figures 1 and S1. Bold numbers indicate best reciprocal BLAST hits.

|  | NHP |  |  | SA | JA |  | GSL |  |
| --- | --- | --- | --- | --- | --- | --- | --- | --- |
|  | ALD1 | SARD4 | FMO1 | ICS1 | OPR3 | JAR1 | SUR2 | SUR1 |
| *Selaginella moellendorffii* | 56 | 0 | 39 | 45 | 53 | 43 | 38 | 41 |
| *Amborella trichopoda* | 63 | 51 | 56 | 52 | 67 | 59 | 44 | 42 |
| *Sesamum indicum* | 69 | 54 | 67 | 58 | 72 | 64 | 49 | 56 |
| *Nicotiana benthamiana* | 65 | 60 | 65 | 50 | 69 | 66 | 50 | 54 |
| *Solanum lycopersicum* | 65 | 58 | 66 | 59 | 74 | 66 | 50 | 56 |
| *Solanum pennellii* | 66 | 58 | 66 | 59 | 74 | 66 | 49 | 55 |
| *Solanum tuberosum* | 65 | 59 | 66 | 59 | 74 | 66 | 47 | 53 |
| *Vitis vinifera* | 70 | 65 | 65 | 63 | 73 | 63 | 50 | 55 |
| *Citrus sinensis* | 70 | 69 | 72 | 61 | 72 | 67 | 50 | 57 |
| *Citrus clementina* | 68 | 69 | 72 | 61 | 53 | 67 | 51 | 57 |
| *Theobroma cacao* | 69 | 68 | 72 | 64 | 75 | 66 | 51 | 56 |
| *Gossypium raimondii* | 70 | 66 | 71 | 63 | 75 | 67 | 48 | 56 |
| *Tarenaya hassleriana* | 78 | 80 | 77 | 75 | 80 | 80 | 90 | 80 |
| *Capsella rubella* | 93 | 95 | 93 | 91 | 93 | 94 | 94 | 92 |
| *Arabidopsis lyrata* | 97 | 95 | 95 | 96 | 95 | 97 | 99 | 98 |
| *Arabidopsis thaliana* | 100 | 100 | 100 | 100 | 100 | 100 | 100 | 100 |
| *Eutrema salsugineum* | 89 | 92 | 91 | 91 | 90 | 91 | 95 | 90 |
| *Brassica napus* | 90 | 89 | 89 | 90 | 90 | 90 | 96 | 89 |
| *Brassica rapa* | 89 | 89 | 89 | 90 | 90 | 90 | 96 | 89 |
| *Eucalyptus grandis* | 64 | 55 | 65 | 62 | 75 | 62 | 45 | 59 |
| *Medicago truncatula* | 69 | 64 | 65 | 54 | 74 | 66 | 47 | 50 |
| *Cicer arietinum* | 68 | 64 | 65 | 55 | 73 | 65 | 45 | 51 |
| *Arachis duranensis* | 67 | 64 | 67 | 56 | 76 | 65 | 49 | 50 |
| *Arachis ipaensis* | 67 | 64 | 67 | 56 | 76 | 66 | 49 | 51 |
| *Lotus japonicus* | 67 | 64 | 28 | 56 | 78 | 62 | 46 | 50 |
| *Phaseolus vulgaris* | 64 | 64 | 67 | 53 | 74 | 64 | 48 | 50 |
| *Glycine max* | 70 | 66 | 67 | 57 | 76 | 65 | 48 | 51 |
| *Vigna radiata* | 68 | 66 | 67 | 55 | 76 | 65 | 47 | 51 |
| *Populus trichocarpa* | 70 | 68 | 69 | 60 | 74 | 68 | 52 | 57 |
| *Ricinus communis* | 68 | 66 | 69 | 62 | 75 | 66 | 51 | 56 |
| *Jatropha curcas* | 69 | 64 | 68 | 60 | 75 | 64 | 51 | 54 |
| *Fragaria vesca* | 68 | 67 | 70 | 61 | 76 | 66 | 43 | 57 |
| *Prunus persica* | 70 | 66 | 70 | 62 | 75 | 66 | 43 | 53 |
| *Prunus mume* | 69 | 66 | 64 | 61 | 76 | 66 | 42 | 52 |
| *Malus domestica* | 68 | 65 | 69 | 62 | 76 | 67 | 43 | 55 |
| *Pyrus x bretschneideri* | 68 | 64 | 69 | 62 | 74 | 65 | 40 | 55 |
| *Cucumis sativus* | 66 | 60 | 66 | 58 | 73 | 67 | 43 | 54 |
| *Cucumis melo* | 66 | 60 | 65 | 58 | 75 | 67 | 44 | 54 |
| *Beta vulgaris* | 69 | 62 | 64 | 58 | 73 | 63 | 49 | 54 |
| *Nelumbo nucifera* | 71 | 58 | 67 | 61 | 72 | 66 | 49 | 57 |
| *Elaeis guineensis* | 65 | 52 | 63 | 55 | 71 | 64 | 44 | 55 |
| *Phoenix dactylifera* | 66 | 54 | 63 | 56 | 70 | 64 | 45 | 57 |
| *Musa acuminata* | 64 | 52 | 60 | 53 | 68 | 63 | 43 | 56 |
| *Oryza sativa* | 64 | 47 | 58 | 53 | 73 | 62 | 39 | 56 |
| *Oryza brachyantha* | 63 | 43 | 60 | 60 | 73 | 61 | 38 | 57 |
| *Brachypodium distachyon* | 63 | 48 | 58 | 55 | 70 | 61 | 39 | 53 |
| *Aegilops tauschii* | 62 | 48 | 59 | 54 | 70 | 61 | 40 | 54 |
| *Zea mays* | 63 | 47 | 59 | 54 | 71 | 62 | 38 | 52 |
| *Sorghum bicolor* | 62 | 47 | 60 | 55 | 71 | 61 | 39 | 53 |
| *Setaria italica* | 64 | 48 | 59 | 53 | 72 | 62 | 43 | 52 |
| *Physcomitrella patens* | 61 | 47 | 29 | 42 | 53 | 41 | 35 | 42 |

